# Force Generation by Enhanced Diffusion in Enzyme-Loaded Vesicles

**DOI:** 10.1101/2025.01.02.627955

**Authors:** Eike Eberhard, Ludwig Burger, César L. Pastrana, Hamid Seyed-Allaei, Giovanni Giunta, Ulrich Gerland

**Affiliations:** Physics of Complex Biosystems, Technical University of Munich, 85748 Garching, Germany

## Abstract

Recent experiments have shown that the diffusion coefficient of some metabolic enzymes increases with the concentration of their cognate substrate, a phenomenon known as enhanced diffusion. In the presence of substrate gradients, enhanced diffusion induces enzymatic drift, resulting in a non-homogeneous enzyme distribution. Here, we study the effects of enhanced diffusion on enzyme-loaded vesicles placed in external substrate gradients using a combination of computer simulations and analytical modeling. We observe that the spatially inhomogeneous enzyme profiles generated by enhanced diffusion result in a pressure gradient across the vesicle, which leads to macroscopically observable effects, such as deformation and self-propulsion of the vesicle. Our analytical model allows us to characterize dependence of the velocity of propulsion on experimentally tunable parameters. The effects predicted by our work provide an avenue for further validation of enhanced diffusion, and might be leveraged for the design of novel synthetic cargo transporters, such as targeted drug delivery systems.

## I. INTRODUCTION

Enzymes have traditionally been viewed as passive components, solely responsible for catalyzing chemical reactions [1]. However, recent findings reveal that enzymes can exhibit properties akin to active matter, in addition to their catalytic activity. For instance, the effective diffusion coefficient of some enzymes has been shown to increase in the presence of its cognate substrate, a phenomenon known as enhanced diffusion [2–6]. While the effect of enhanced diffusion has been validated through a range of different experimental assays [7–10], the microscopic mechanism underlying this effect remains under debate [11–15]: some attribute enhanced diffusion to non-propulsive mechanisms such as conformational changes [8, 16], whereas others emphasize the role of dissipative propulsion [10], for instance, due to phoresis [2, 3] or pressure waves [5]. Regardless of the microscopic mechanism, enhanced diffusion results in the directed motion of enzymes in the presence of a substrate gradient. It causes enzymes to drift from regions of high diffusivity (high substrate concentration) towards regions of low diffusivity (low substrate concentration), resulting in a non-homogeneous enzyme density [6, 7].

In this work, we propose a micro-device that uses forces generated by enzymatic enhanced diffusion for its own deformation and propulsion. Using coarse-grained simulations and analytical modeling, we demonstrate that enzymes encapsulated in a micron-sized vesicle can exert forces of a few piconewtons when they exhibit enhanced diffusion in a substrate gradient. These forces cause shape deformations, alter the fluctuation spectrum, and propel the vesicle with appreciable velocities. Motile micro- and nano-devices hold significant potential for practical applications, particularly in the design of synthetic cargo transporters [17, 18], such as biocompatible drug delivery systems [19–21]. Existing systems typically achieve motion through slip velocities generated by self-induced gradients [17]. Such gradients can arise, for example, from an inhomogeneous enzyme distribution on the surface of a moving bead [22–24] or vesicle [25], or from an inhomogeneous permeability of a vesicle to product molecules produced inside [19]. The mechanism that causes the motion in our device is different: Enzymes encapsulated within the vesicle form a non-homogeneous profile, creating a difference in osmotic pressure across the vesicle, which drives its deformation and motion. Unlike existing systems, our micro-device does not need to be inherently asymmetric: The asymmetry emerges dynamically via enzymatic enhanced diffusion due to a pre-imposed external substrate gradient.

Our results suggest that the morphological changes and the motion expected for our micro-device should be detectable by light microscopy, providing an avenue for further validation of enhanced diffusion. Moreover, our analytical model rationalizes the dependence of the device’s behavior on enzyme properties and experimentally tunable parameters. This model can guide the design of synthetic bio-compatible cargo systems with controllable active properties.

## II. RESULTS

### A. System Description

We consider a vesicle containing enzymes *ε* with cognate substrate *s*. The vesicle is submerged in an external substrate gradient, and it is assumed to be perfectly permeable to substrate and impermeable to enzymes (Fig. 1). This assumption is justified by the (typically) much larger size of enzymes compared to that of their substrate. We neglect substrate depletion, such that the internal concentration of *s* is set by the external substrate gradient. A rapid permeation of substrate can be achieved by incorporating natural or artificial pores on the membrane [26, 27] (SI Sec. I). We further assume that the enzymes undergo enhanced diffusive motion (Fig. 1, inset), where the diffusivity of the enzymes is a function of the substrate concentration *s*(**r**) at position **r** [14],

**Figure 1.**
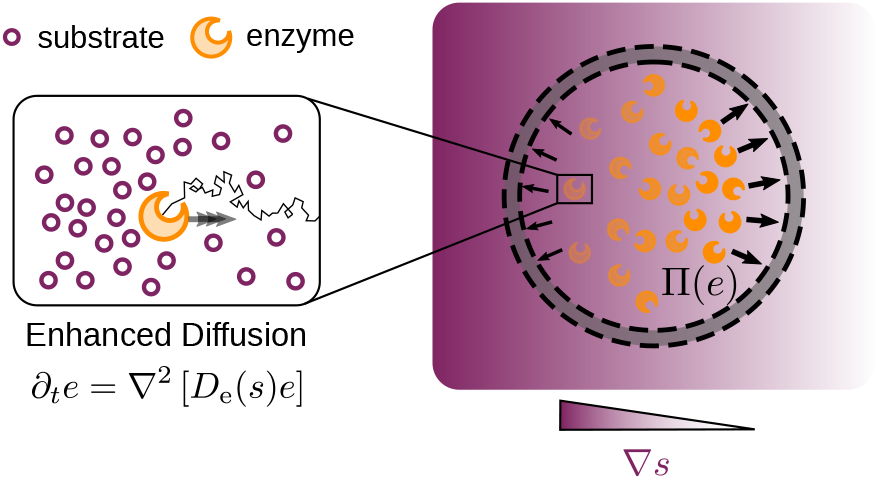
Mechanism generating anisotropic pressure profile. A vesicle is loaded with enzymes that catalyze the conversion of substrate *s*. While the substrate is assumed to freely permeate through the membrane, enzymes cannot pass the membrane. The vesicle is placed in an externally imposed substrate gradient ∇*s*, depicted in the figure by a purple shading. *Inset:* The time-evolution of the enzyme is governed by an enhanced diffusion equation, i.e., the enzyme diffusion *D*_e_ is a function of the substrate concentration *s*. Enhanced diffusion causes the enzyme to move downstream substrate gradients (towards low substrate concentration). This results in a non-homogeneous enzyme distribution within the vesicle and leads to an anisotropic pressure profile Π(**r**, *t*).

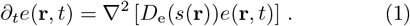

The effective diffusion coefficient *D*_e_(*s*) is defined as [5, 7, 28]

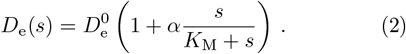

The factor α parameterizes the relative increase in diffusivity 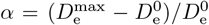 due to the substrate, with typical values in the range of 15-80% and an average of approximately 30% [13].

In the presence of the substrate gradient, enhanced diffusion (Eq. (1)) results in antichemotactic motion of the enzymes, which causes the accumulation of enzymes in regions of low substrate concentration [6, 7]. We note that other studies report motion towards regions of high substrate concentration (chemotaxis) [2, 28, 29], which can be attributed to cross-diffusion contributions due to the (non)specific interactions of enzymes with substrate and product molecules [30, 31]. However, cross-diffusion effects have been shown to dominate for high substrate/product concentrations (*s* ≫ *K*_M_), and can be neglected in our model. The spatially inhomogeneous enzyme profile created by antichemotaxis (Eq. 1) creates an anisotropic osmotic pressure on the membrane, Π(**r**, *t*) = *K*_B_*T e*(**r**, *t*), with *T* the temperature of the solvent [32, 33]. The pressure profile exerts a net force onto the vesicle,

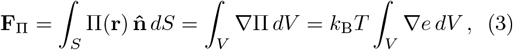

where 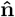(**r**) is the outward normal vector of the surface *S*, and *V* is the volume of the vesicle.

We estimate the magnitude of the force for a simplified scenario: (i) The substrate concentration is assumed to be a linear function of *x*, implying constant ∇*s*, (ii) the enzyme equilibration timescale is assumed to be small compared to the timescale associated with any possible deformation/displacement of the vesicle, and (iii) the vesicle is assumed to retain spherical shape. Using the steady state solution of Eq. (1), we obtain an equation for the force exerted by the enzymes (Eq. (24) in the Methods Section). Enzymes with the properties of urease (*α* = 0.24, *K*_M_ = 3 mM, and 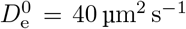) that are encapsulated in a vesicle of size *R* = 10 µm at a concentration of *e*_T_ = 100 nM and placed in a substrate gradient, ∇*s* = 0.05 mM/µm, exert a force of about 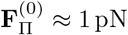. This force is in a biologically relevant range [34].

### B. Enzymes undergoing enhanced diffusion can deform vesicles

As a first step, we aim to probe the deformation of the vesicle due to the pressure exerted by the enzymes. To this end, we simulate the dynamics of the system depicted in Fig. 1 using a well-established model that is known to capture the equilibrium fluctuation spectrum of lipid membranes and their deformation induced by active particles correctly [35–38]. The model accounts for bending elasticity, membrane fluidity, area conservation, and a volume constraint (Methods section). To effectively describe the local pressure profile, Π(**r**, *t*), we introduce a repulsive short-range interaction between the vesicle surface and the enzyme particles. That way, enzymes exert an effective force onto the membrane, proportional to their local concentration. The motion of enzymes is described by an overdamped Langevin equation with multiplicative noise. Using Itô’s interpretation of stochastic calculus [39], it can be shown that this Langevin equation is equivalent to the enhanced diffusion equation Eq. (1) (see Methods Section and SI Sec. II).

Our simulations indicate that enzymes encapsulated in vesicles cause substantial deformations as a consequence of enhanced diffusion (Fig. 2). However, the magnitude of the deformation depends on the osmolarity of the solution that surrounds the vesicle, i.e., the concentration of osmotically active solutes outside of the vesicle that exert osmotic pressure in addition to the enzymes. To account for the osmolarity of the solution, we incorporate a volume constraint into the vesicle model, which allows us to study the system under hyperosmotic (high concentration of osmotically active solutes) and hypoosmotic conditions (low concentration of osmotically active solutes) conditions [40].

**Figure 2.**
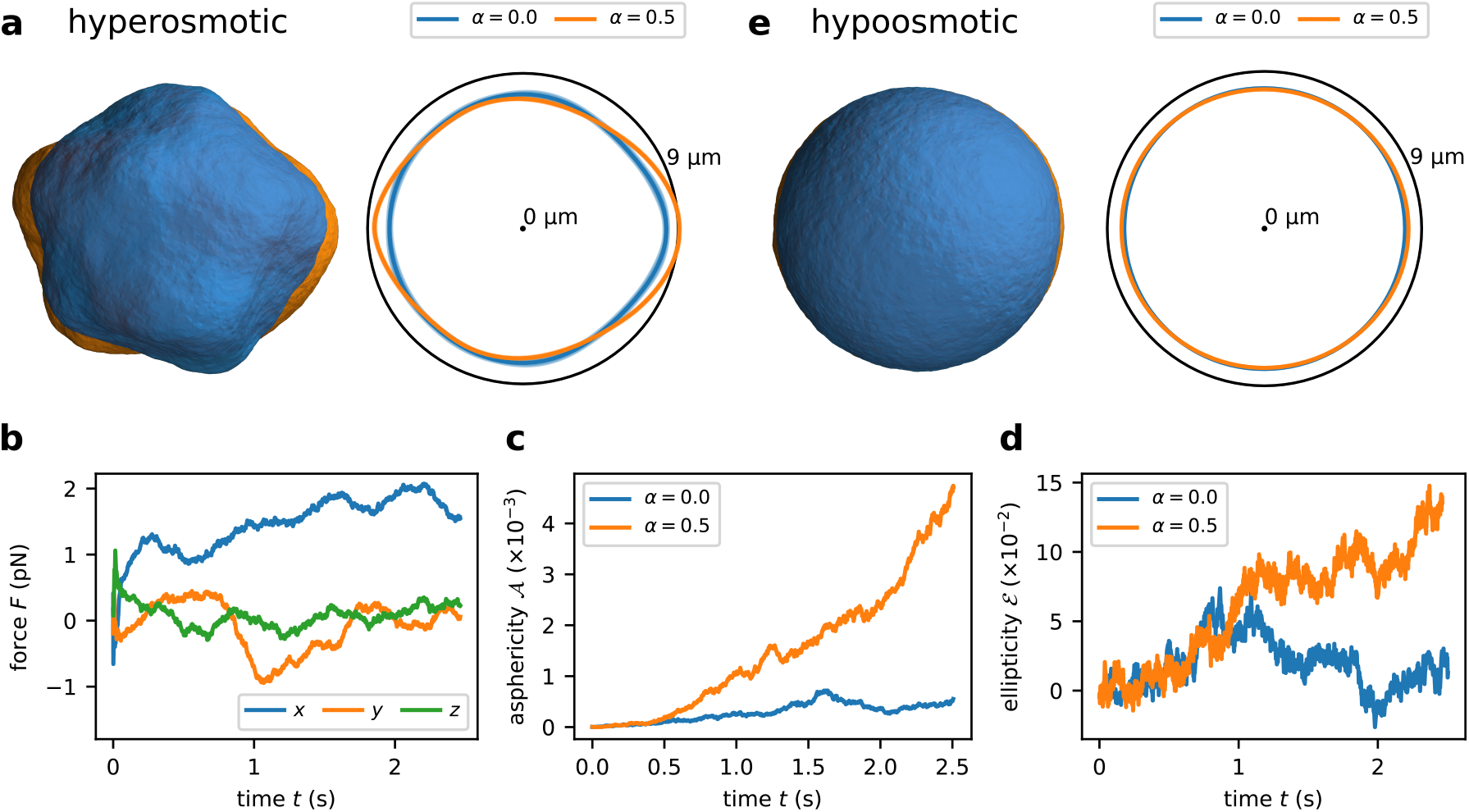
Deformations of enzyme-loaded vesicles. a)-d) Vesicles under hyperosmotic conditions (*K*_*V*_ = 129 N m^−2^, 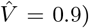 filled with enzymes that do (*α* = 0.5, orange) or do not exhibit enhanced diffusion (*α* = 0.0, blue) in the presence of an external substrate gradient. a) Snapshots of the vesicle shape (*left*) and averaged cross sections (*right*) show a prolate deformation of the vesicle due to enhanced diffusion. The averaged cross-sections are obtained by averaging over multiple time frames and rotations along the gradient axis: The continuous lines represent the local average radius, while the standard deviation is shown as shaded area. b) Components of the force vector acting on vesicles under hyperosmotic conditions as a result of enzyme-vesicle interaction. The force component parallel to the direction of the gradient is non-zero. c)-d) The shape parameters aspheriticy 𝒜 and ellipticity *ε* in vesicles under hyperosmotic conditions are non-zero, indicating a prolate deformation of the vesicle. e) Shape of vesicles under hypoosmotic conditions (no volume constraint) filled with enzymes that do (*α* = 0.5, orange) or do not exhibit enhanced diffusion (*α* = 0.0, blue) in the presence of an external substrate gradient. Snapshots (*left*), averaged cross-sections (*right*) and shape parameters (Fig. S2) indicate a subtle prolate deformation due to enhanced diffusion. The parameters used in the simulations are summarized in SI Table II.

Under hyperosmotic conditions, the vesicle shows clear deformations (including protrusions) even in the absence of enhanced diffusion (Fig. 2a, left). Since only a twodimensional projection of the three-dimensional shape can be observed in a typical light microscopy setup, we determine the shape of this projection by averaging over the different cross-sections obtained through rotation around the gradient axis. The cross-section indicates a preferential stretching of the vesicle along the direction of the substrate gradient for *α* = 0.5 (Fig. 2a, right). The force generated along the *x*-axis in the presence of enhanced diffusion (*α* = 0.5) builds up, reaching values ≈ 2pN after 2s (Fig. 2b), which is comparable to the estimate we obtained previously.

To further characterize the shape of vesicles under hyperosmotic conditions, we resort to a set of widely employed order parameters (see SI Sec. II D and [41–43]). For example, the asphericity 𝒜 measures the deviation of the shape from a perfect sphere. For *α* = 0.5, we observe a continuous increase of 𝒜, reaching values more than 5 times larger than in the absence of enhanced diffusion (Fig. 2c). Similarly, the ellipticity along the equatorial plane *ε* is also 3 to 5 times larger for vesicles loaded with enzymes that exhibit enhanced diffusion (Fig. 2d). Interestingly, both 𝒜 and *ε* do not saturate within the simulated time-window, indicating that larger deformations can be expected at longer times.

Under hypoosmotic conditions (Fig. 2e), vesicles are mostly spherical and exhibit only slight deformations. Although the deformation is hard to detect by a simple visual inspection of the vesicle shapes, a closer analysis of the cross-section reveals a subtle prolateness along the gradient axis. This is consistent with an ellipticity of *ε* ∼ 2–3% (Fig. S2). The vesicle reaches a steady-state shape after approximately 1s, as indicated by the shape observables approaching a constant asymptotic value.

Inspired by recent works focused on vesicles loaded with active particles [38, 43], we aimed to determine if the fluctuation spectrum of the vesicle’s radius reveals signatures of deformation (Methods section). The analysis of membrane fluctuations is a well-established experimental technique to quantify the shape and mechanics of lipid membranes [44–47]. We find that the fluctuation amplitudes *a*_𝓁_ in the absence of enhanced diffusion *α* = 0 coincide perfectly with the theoretical prediction derived for fluctuating membranes in equilibrium with a surface tension of σ = 0.97 µN m^−1^ (Fig. 3a, see Methods Section and SI Sec. III). The fluctuation spectrum in the presence of enhanced diffusion *α* = 0.5 exhibits significant deviations for the lowest modes 𝓁 = 2 and 𝓁 = 3. These modes correspond to elliptic and triangular deformations (Fig. 3b), consistent with the shape observed in Fig. 2e.

**Figure 3.**
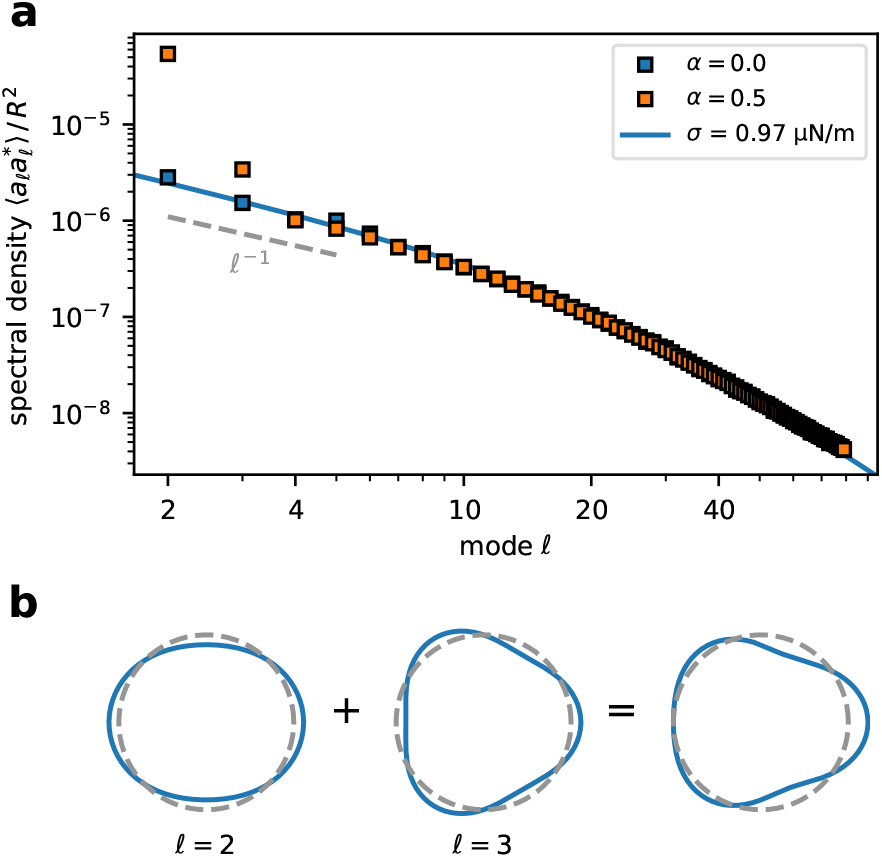
Morphology of enzyme-loaded vesicles. a) Fluctuation spectra of vesicles under hypoosmotic conditions (no volume constraint) filled with enzymes that do (*α* = 0.0, red) or do not display enhanced diffusion (*α* = 0.5, blue). The second and the third fluctuation modes are significantly more pronounced for *α* = 0.5 than for *α* = 0.0. The characteristic scaling in the tension-dominated regime *a*_𝓁_ ∼ 𝓁^−1^ is highlighted as dashed line. The continuous blue line is the spectral density predicted by the model described in SI Sec. III. The parameters used in the simulations are summarized in SI Table II. b) Schematic illustration of the deformations corresponding to the two fluctuation modes that are boosted by enhanced diffusion.

### C. Enzymes undergoing enhanced diffusion can propel vesicles

In addition to the deformation, we observe that the vesicle propels itself through the externally imposed gradient under hypoosmotic conditions. We introduce a simplified model where the vesicle is treated as a perfectly non-deformable sphere subjected to Stokes’ friction (Methods Section). This model is applicable since deformations have been shown to be small under hypoosmotic conditions. Unlike the mesh-based model used before, the simplified simulation accurately captures vesicle friction (SI Sec. II E), predicting quantitatively correct propulsion velocities. The center of mass of the vesicles moves toward regions of lower substrate concentration. The vesicle translates at a velocity 𝑣 = 0.61 µm s^−1^ (Fig. 4), which is a factor ∼ 70 slower than the velocity expected based on the steady-state enzyme distribution *e*(*r*) (obtained by solving Eq. (1) in steady state) and its associated force on the vesicle **F**_Π_ (Eq. (3)). Interestingly, the enzyme profile observed in the simulation is more homogeneous in space (Fig. 4b) than the steadystate distribution predicted by Eq. (1), implying that the vesicle motion (and the associated drift of enzymes) alters the enyzme profile. In control simulations in which the location of the vesicle is constrained to the origin, the force and the enzyme profile are in excellent agreement with the steady-state solution of Eq. (1) (Fig. S4). This suggests that vesicle motion needs to be accounted for to predict the enzyme profile and the forces exerted on the membrane accurately.

**Figure 4.**
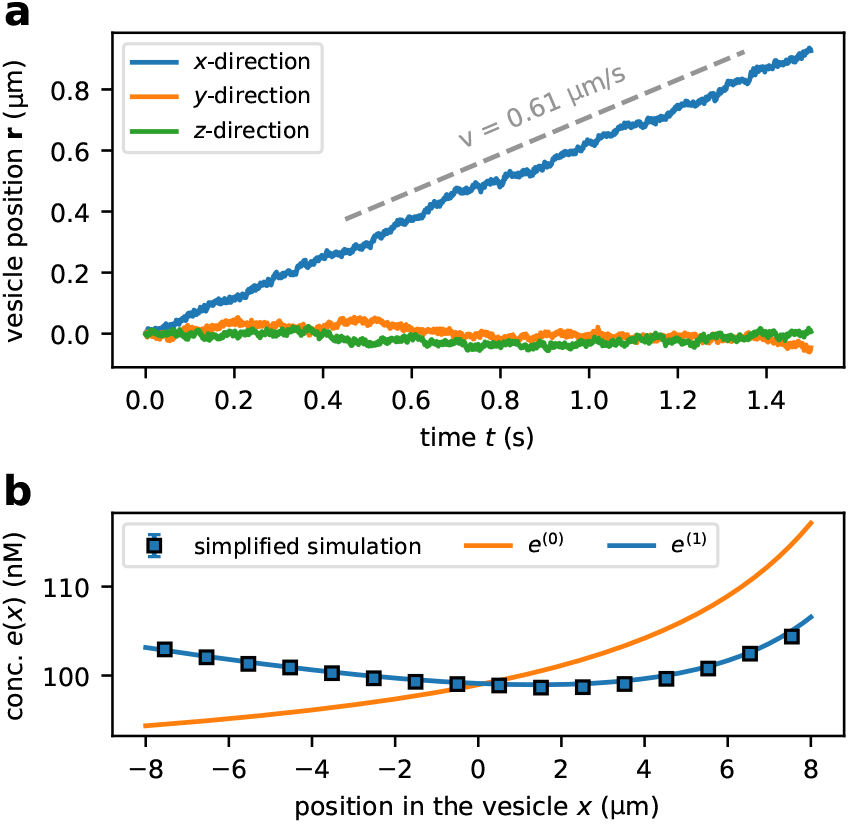
Self-propulsion of enzyme-loaded vesicles. a) Time-evolution of the vesicle position (center of mass) obtained via the simplified simulation. The vesicle moves downstream along the substrate gradient with velocity 𝑣 determined via a linear fit. Vesicle trajectories obtained via the full mesh-based simulation are shown in Fig. S3. b) Enzyme concentration profile along the gradient axis. The profile obtained to linear order in Péclet number, *e*^(1)^, agrees well with the profile observed in the simplified simulation. This profile is more homogeneous than the adiabatic profile *e*^(0)^ (steadystate solution of Eq. (1)). The dots represent the average and the standard error of the mean of at least *n* = 10 simulations. The uncertainties are smaller than the label size. The parameters used in the simulation are summarized in SI Table III.

Hence, we describe enzyme diffusion in the co-moving frame, where the vesicle is at rest by including an additional drift term in Eq. (1),

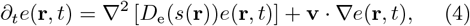

where **v** is the velocity of the vesicle. Under the assumption that the vesicle velocity is only non-zero along the direction of the substrate gradient, Eq. (4) simplifies to a one-dimensional differential equation. The resulting steady state enzyme profile is a function of the dimensionless Péclet number 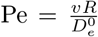, which relates the timescale of diffusion to the timescale of vesicle drift. Even at the highest velocities observed in the simulation, 𝑣 ≈ 0.6 µm s^−1^, for a vesicle with radius *R* = 8 µm and diffusion coefficient 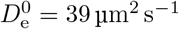, the Péclet number is small, Pe ≈ 0.12. This allows us to expand the full steady state enzyme profile in the Péclet number (see Methods Section and SI Sec. IV).

To zeroth order (only terms independent of the Péclet number), we recover the steady-state profile and velocity obtained by solving Eq. (1), 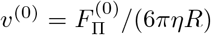 (Methods Section). As neglecting contributions proportional to the Péclet number is equivalent to assuming that the translation of the vesicle is much slower than the enzyme diffusion, we refer to the limit Pe *→* 0 as the adiabatic limit. The steady state velocity to linear order in Péclet number 𝑣 ^(1)^ can be expressed in terms of the adiabatic velocity 𝑣 ^(0)^,

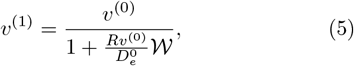

where *𝒲* is a non-negative weight that depends on the dimensionless position-dependent diffusion coefficient 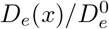 as well as on the vesicle radius *R*. From Eq. (5), we can directly see that the translation velocity is well-approximated by the adiabatic velocity,

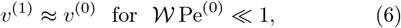

provided the Péclet number associated with the adiabatic velocity, 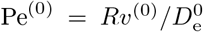, is small. In the opposite limit, the translation velocity approaches an asymptotic value that is independent of the adiabatic velocity,

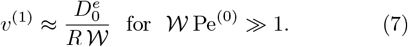

In both limits, the translation velocity of the vesicle depends on its position in the system. As the vesicle moves, the substrate concentration within the vesicle changes, which affects the effective diffusion coefficient of the enzymes, and, consequently, the pressure and the ensuing translation velocity. Therefore, the velocity observed in the simulation corresponds to an effective velocity, averaged over the positions of the vesicle along its trajectory. Aiming for a direct comparison of the theoretical prediction to the simulations, we analytically compute an effective velocity 𝑣 based on 𝑣 ^(1)^ by averaging 𝑣 ^(1)^(*x*) over the vesicle’s trajectory (Methods Section). The analytical theory agrees well with the simulation data with regard to the predicted enzyme profile (Fig. 4b) and translation velocities (Fig. 5).

**Figure 5.**
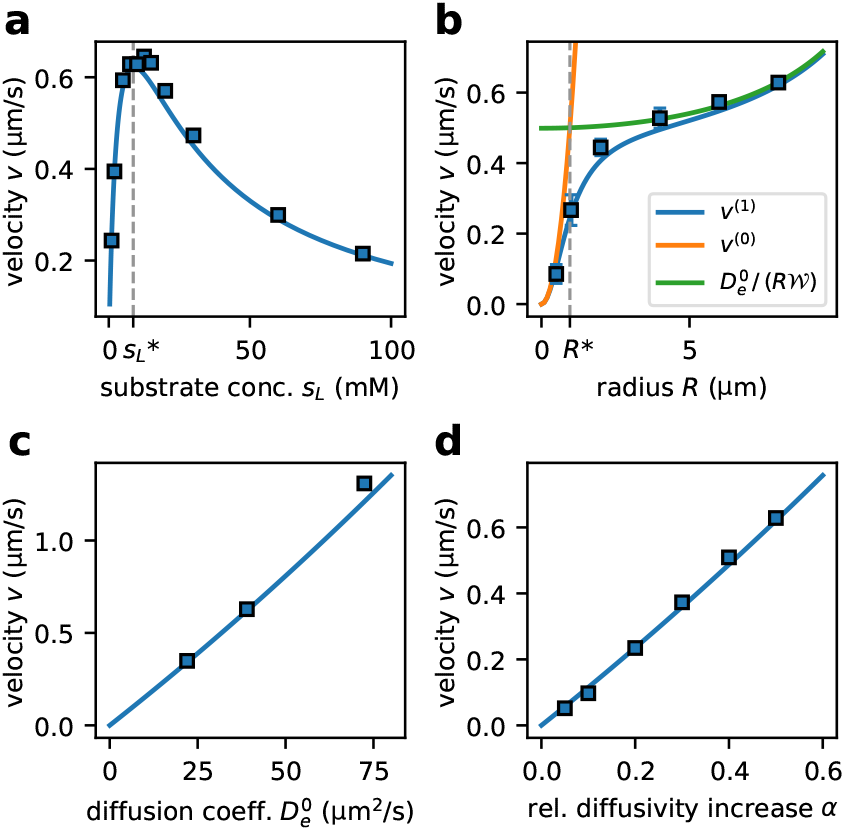
Dependence of vesicle velocity on system parameters and enzymatic properties. a) Vesicle velocity 𝑣 depends non-monotonously on the substrate gradients, and exhibits a maximum at intermediate substrate concentration 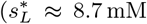, shown as dashed line). b) Velocity increases as a function of vesicle radius, with a quadratic radius-dependence for sufficiently small radii (smaller than *R*^*\**^ ≈ 0.96 µm, shown as dashed line). c) Velocity increases linearly with the enzymatic basal diffusion coefficient 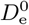. d) Relation between the enhanced diffusion factor *α* and the velocity. All panels show the translation velocity in a medium with viscosity similar to that of water, 𝜂 = 1 mPa s (parameters are summarized in SI Table III). The dots represent the average of at least *n* = 10 simulations. The standard deviation of the mean is usually smaller than the marker size. The continuous curves show the effective trajectory-averaged velocity computed via the self-consistency approach to linear order in Péclet number 𝑣 ^(1)^. Fig. S11 shows the parameter-dependence of velocity in a medium with high viscosity under otherwise identical conditions.

In Fig. 5a, we show the velocity 𝑣 as a function of the height of the substrate gradient *s*_l_. The velocity depends non-monotonically on the substrate concentration, with a peak in velocity at around *s*_l_ ≈ 10mM. This behavior of the velocity is a consequence of the substrate-dependence of the diffusion coefficient *De*(*x*). In the linear regime of the Michaelis-Menten response curve (*s* ≪ *K*_M_), the diffusion coefficient depends linearly on the substrate concentration. Thus, a steep substrate gradient enhances the difference in enzyme diffusivity between the left and right sides of the vesicle, leading to a more pronounced gradient in enzyme concentration, and to higher translation velocities. At saturating conditions (*s* ≫ *K*_M_), the diffusion coefficient is constant everywhere within the vesicle, which causes the pressure difference between left and right end of the vesicle (and consequently the translation velocity) to vanish. Solving for the substrate concentration 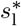 at which the velocity attains its maximum reveals that 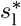 depends linearly on *K*_M_,

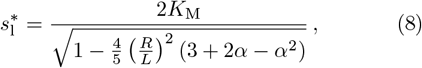

see dashed line in Fig. 5a and SI Sec. IV D. We note that 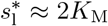 for vesicles that are far smaller than the system size, *R* ≪ *L*.

The radius-dependence of the velocity is shown in Fig. 5b. Intuitively, the force *F* exerted by the enzymes is expected to be proportional to the number of enzymes in the vesicle, such that *F* ∝ *R*^3^ for fixed enzyme concentration. Together with Stokes’ law, the scaling of the force implies that the velocity should depend quadratically on the radius, 𝑣 ∝ *R*^2^. This intuition is consistent with the quadratic radius-dependence of the adiabatic velocity 𝑣^(0)^ (SI Sec. IV E). As long as the vesicle radius is small enough for the adiabatic Péclet number Pe^(0)^ to be small, the translation velocity to be dominated by the adiabatic velocity, and the velocity scales quadratically with *R* (see orange curve in Fig. 5b). However, as the radius increases, the adiabatic Péclet number Pe^(0)^ is not small anymore, and corrections beyond the adiabatic limit contribute significantly to the translation velocity. The characteristic radius *R*^*^ beyond which corrections to the adiabatic limit are non-negligible is set by the condition *𝒲*(*R*^*^) Pe^(0)^(*R*^*^) = 1,

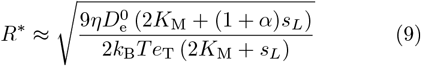

(see vertical dashed line in Fig. 5b). In the limit of large radii, *R* ≫ *R*^*^, the translation velocity is given by the radius-dependent weight *𝒲* (Eq. (7)), leading to 𝑣 ∼ (*c*_0_ + *c*_1_*R*^2^ + *c*_2_*R*^4^)^−1^, where *c*_*i*_ are radius-independent constants (see green curve in Fig. 5b, and SI Sec. IV E).

Besides the aforementioned system parameters (strength of the substrate gradient, vesicle size), enzyme properties, such as the enzymatic diffusion coefficient 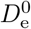, also affect the translation velocity. We simulate the vesicle translation considering the diffusion coefficients of urease, hexokinase and acetylcholinesterase, all of which are enzymes for which experimental evidence of enhanced diffusion exists [6, 28]. We find that the velocity depends linearly on the basal diffusion coefficient. This is a consequence of Pe^(0)^ ≪ 1 over the entire range of diffusion coefficients (SI Sec. IV F):

The translation velocity is given by Eq. (7), and thus proportional to 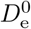. We also investigate the effect of *α* on the velocity (Fig. 5d). Our results indicate that 𝑣 is directly proportional α. This can be rationalized by noting that increasing *α* (and thus the effective enzymatic diffusion coefficient) leads to a decrease of the diffusion timescale in a similar way as increasing 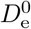 does.

## III. CONCLUSIONS AND DISCUSSION

Over the last years, numerous studies have established [3–6] and validated [7, 9, 10] that certain enzymes exhibit enhanced diffusion in the presence of their cognate substrate. In this work, we have shown that enzymatic enhanced diffusion can be leveraged to design micron-sized devices capable of self-propulsion: Enzymes are encapsulated within a vesicle (Fig. 1), and establish a spatially inhomogeneous enzyme profile due to an antichemotactic drift caused by enhanced diffusion (Fig. 4b). This enzyme profile generates a gradient of osmotic pressure across the vesicle, driving its deformation (Fig. 2 and Fig. 3) and propulsion (Fig. 4 and Fig. 5). While the prolate deformation of the vesicle is clearly visible under hyperosmotic conditions (Fig. 2a-d), deformations are less pronounced in hypoosmotic conditions (Fig. 2e). The vesicle propels itself with velocites on the order of 1 µm s^−1^ (Fig. 4). We have characterized the parameter-dependence of the velocity using an analytical model (Fig. 5): The velocity depends linearly on the bare enzymatic diffusion coefficient, 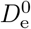, or the strength of enhanced diffusion, *α* (Fig. 5c-d), and it increases non-linearly with vesicle radius, *R*, or substrate gradient (Fig. 5a-b). Substrate concentrations exceeding *K*_M_ can even decrease the propulsion velocity due to enzyme saturation.

In our model, we focused on the propulsion velocity arising from the gradient in osmotic pressure driven by enhanced diffusion, but neglected hydrodynamic effects, which can provide an additional contribution to the propulsion velocity: Gradients in the concentrations of substrate, product or enzyme can cause nonzero slip velocities via phoretic effects [24, 48–50] or through a concentration-dependent line tension of the vesicle (Marangoni flow) [17, 50, 51], generating flow fields within and/or outside of the vesicle. The magnitude of these additional velocity contributions depends on the details of the experimental setup, such as the properties of substrate and product (e.g., their charge or pK_a_), and the properties of the encapsulating vesicle (e.g., its line tension or size).

Recent advances in synthetic biology open up the opportunity to implement the micro-device proposed in this work in laboratory experiments: Vesicles of sufficient size (up to 100 µm in radius) can be prepared [52], and, once loaded with enzymes, placed in a substrate gradient produced, e.g., by a microfluidic device. Unlike nano-devices that rely on their intrinsic asymmetry for their function [19, 23, 24], enzyme-loaded vesicles driven by enhanced diffusion do not require any asymmetry to be introduced during the fabrication process. Instead, the asymmetry is dynamically generated by enhanced diffusion in the presence of a substrate gradient. We suppose that this property simplifies the fabrication of the device. Furthermore, the control of its motion should be simpler compared to nano-devices, for which the direction of propulsion depends sensitively on potentially hard to control parameters, such as the surface composition [19, 23] and shape [53]. In our case, the vesicle always moves towards regions of low substrate concentrations (antichemotaxis) as a consequence of the antichemotactic drift of enhanced diffusion.

Motile nano- and micro-devices exhibiting (anti)chemotaxis are of great practical importance for the design of synthetic cargo transporters [17], such as biocompatible drug delivery systems [19–21]. For example, the ability of nanoswimmers to penetrate the blood-brain barrier has been shown to be enhanced by chemotaxis in response to a gradient of glucose [19]. Similarly, antichemotaxis in response to a gradient in oxygen (oxygen decreasing along the depth of the tumoral tissue) might guide a swimmer towards the center of a tumor [54]. Further research is needed to determine the experimental feasibility and practical applicability of the enhanced-diffusion based microswimmer. Its characterization provided in this work may help to navigate the vast space of tuneable system parameters, and aid the rational design of such a device.

## METHODS

### Mesh-based Vesicle Model

The vesicle is simulated using a coarse-grained dynamically triangulated mesh consisting of *N*_*V*_ vertices [38] (SI Sec. II A). The Helfrich bending energy is described by the well-known equation *U*_*b*_ = 2*k ∬ H*^2^d*S*, where *H* is the mean curvature, *κ* is the bending modulus [55]. This energy is discretized on the mesh [56],

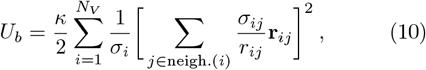

where the sum runs over all vertices *N*_*V*_ and **r**_*ij*_ vector from node *j* to node *i*. The term σ_*ij*_ corresponds to the length of the bond in the dual lattice 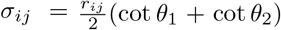 where *θ*_1_ and *θ*_2_ are the angles opposite to the bond *i*-*j*. The term σ_*i*_ is a normalization factor and accounts for the total area of the dual cell around vertex *i* and is given by 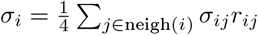.

The area of the vesicle membrane is fixed by harmonic potential acting locally on the triangles of the mesh [38,42]:

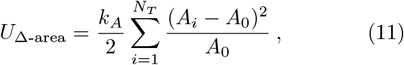

where *A*_*i*_ are the instantaneous triangle areas, 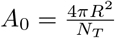 and *K*_*A*_ is the area conservation coefficient. The sum runs over all triangles *N*_*T*_ = 2(*N*_*V*_ − 2) [38, 42]. To account for osmotic effects [40], when specified, we use the harmonic potential

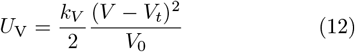

to constrain the volume, where *K*_*V*_ is the volume stiffness and *V*_*t*_ is the target volume of the vesicle. We define 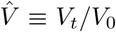, where 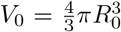. The volume enclosed by the non-convex polyhedron is determined as described in [57, 58].

In the absence of volume constraints, the pressure exerted by the enzymes (both with and without enhanced diffusion) inflates the vesicle, stretching the surface: the vesicle is in a high-tension state. Conversely, introducing a volume constraint mimics hyper-osmotic conditions (by setting a target volume 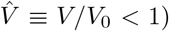, which acts against the pressure exerted by the enzymes and the vesicle is at low-tension conditions.

The stability of the mesh is maintained by the bond potential

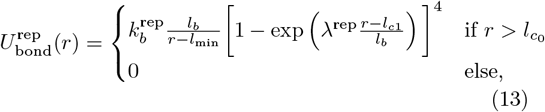

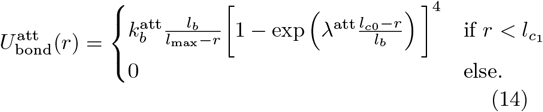

Here, 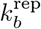 and 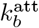 are the bond stiffness parameters. The values *l*_min_ and *l*_max_ set the minimum and maximum bond lengths allowed, and the constants λ^rep^ and λ^att^ control how rapidly the potentials increase between the cutoff and the limiting bond lengths. The parameters *l*_*c*0_ and *l*_*c*1_ are the cutoff lengths of the attractive and repulsive term of the bond potential, respectively, and thus, membrane vertices can move freely in the range 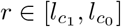.

To account for the fluidity of lipid bilayers, we allow the triangulation to change dynamically [42, 59]. In this procedure, the edges shared by adjacent triangles can be flipped. If a flip occurs, the bond along the shared edge is replaced by a bond connecting the previously unconnected vertices. A randomly selected fraction *ψ* of all bonds is tested at a frequency of *ω*. The parameters *ω* and *ψ* define the viscosity of the membrane [42], and we employed the values described in [38]. To preserve the stability of the mesh, we discard from bond flipping vertices of the shared edge with coordination number ≤ 5. Finally, a bond flip trial is accepted or rejected following the Metropolis criterion with probability 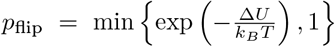, where_flip_ Δ*U* = *U*_flipped_ − *U*_not flipped_ ≈ Δ*U*_bond_ + Δ*U*_Δ-area_. Enzymes are described as particles that do not interact with each other. The interaction between enzymes and the vertices of the membrane is harmonic and of the form

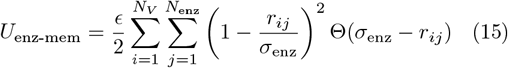

Here *r*_*ij*_ = ∥ **r**_mb,*i*_ − **r**_enz,*j*_ ∥ is the distance between the coordinates of the enzymes and the membrane vertices and the sum runs over all the *N*_*V*_ vertices and all the *N*_enz_ pairs. The Heavyside function Θ restrains the potential to distances *r* < σ_enz_.

### Simplified Vesicle Model

In the simplified simulation, the vesicle is modeled as a non-deformable sphere centered. The vesicle is uniquely defined by its center **r**_mem_ (which evolves in time) and its (time-independent) radius, and there is no need to represent the vesicle membrane as a mesh of vertices. The vesicle membrane interacts with the enzymes via the potential,

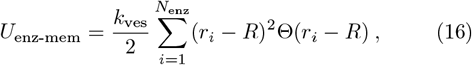

where *r*_*i*_ = ∥**r**_enz,*i*_ − **r**_mem_∥ and *K*_ves_ is the vesicle stiffness. This potential confines all the enzymes within a spherical volume of radius *R*.

### Dynamics of the System

For both vesicle models (mesh-based and simplified), we model the motion of enzymes by an overdamped Langevin approach (Brownian Dynamics),

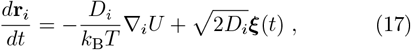

where **r**_*i*_ are the coordinates of the particle *i* (SI Sec. II B). The equation accounts for the random forces due to the collisions with the surrounding fluid (modeled as Gaussian white noise), as well as for a drift due to the potential *U*. Here, the relevant potential is the membrane-enzyme interaction potential *U*_enz−mem_. The diffusion coefficient of the enzymes is a function of the local substrate concentration and, consequently, a function of space *D*_e_ = *D*_e_(*s*(**r**_e_)) (Eq. 2). We consider a linear substrate profile of the form

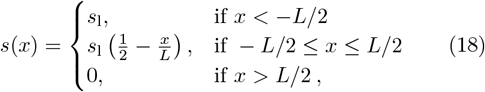

with the vesicle initially placed at the origin *x* = 0. The length of the system *L* is chosen to be 20% longer than the largest simulated vesicle.

In the mesh-based vesicle model, the vertices follow the same Langevin equation as the enzymes. The vertex diffusion coefficient is different from the enzyme diffusion coefficient (see [38] and SI Table II) and a potential *U* that includes all contributions to the membrane energy listed in the explanation of the mesh-based vesicle model. In the simplified vesicle model, there are no vertices to represent the membrane. Instead, the center of mass follows the Langevin equation Eq. (17) with a diffusion coefficient set by the radius of the vesicle and the viscosity of the surrounding medium, *D*_mem_ = *K*_B_*T*/(6*π𝜂R*). In all models, the equations of motion are integrated using the first-order Euler-Maruyama method. The initial enzyme distribution is chosen to be homogeneous or to follow the adiabatic enzyme profile *e*^(0)^ (steady state solution of Eq. 1) depending on the context (SI Sec. II C).

### Fluctuation Analysis

We characterize the morphology of the vesicle via shape parameters (SI Sec. II D) as well as via its fluctuation spectrum. To determine the latter, we conduct a Fourier analysis of the radial fluctuations of an equatorial cross-section of the vesicle [38, 44]. Vertices within a distance on the order of the bond length of the mesh *l*_*b*_ to the equatorial plane are orthogonally projected on that plane. The resulting 2D coordinates are transformed to polar coordinates (*x, y*) *→* (*r, θ*), and linear interpolation is used to obtain *N* radial positions *r* at evenly spaced intervals for *θ*. The resulting radial positions are decomposed in Fourier modes,

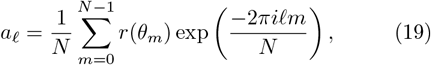

where *θ*_*m*_ = 2*πm*/*N* and 𝓁 is the mode number. We obtain the mean fluctuation spectrum by averaging spectra over *N*_*ρ*_ rotations and over a time window given by *N*_*s*_ uncorrelated frames,

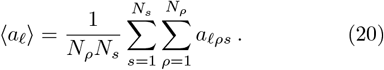

The rotated configurations are computed using a linear transformation of the coordinates of the membrane vertices **r**_mem_, following **r**^′^_mem_ = ℛ_*x*_**r**_mem_, where *ℛ*_*x*_ = *ℛ*_*x*_(*α*) is the rotation matrix along the substrate gradient axis by an angle *α ∈* [0, *π*). Note that a similar approach was used for the cross-section representations shown in Fig. 2a/e.

For a system in equilibrium, obeying the equipartition theorem, the fluctuations 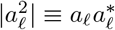 are given by [60]

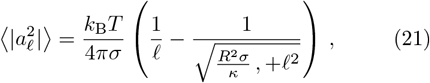

where σ is the surface tension and *κ* is the bending stiffness of the membrane; both parameters can be computed based on the membrane Hamiltonian (SI Sec. III). Eq. (21) approches 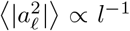 in the so-called tension σ*k*_−1_ ≫ *l*^2^*R*^−2^ and 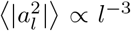 in the bending dominated regime (σ*κ*^−1^ ≪ *l*^2^*R*^−2^).

The illustrations in Fig. 3b are obtained via the inverse Fourier transform considering the following identity for a real-valued signal 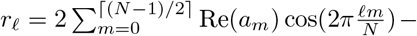 Im(*a*_*m*_) sin 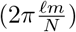. The zeroth mode 𝓁 = 0, which captures the mean radius, and the first mode 𝓁 = 1, which parameterizes translation-like fluctuations, are not relevant in this context.

### Translation Velocity via Self-Consistency Approach

The enzyme profile obtained by finding the steadystate solution of Eq. (4) reads

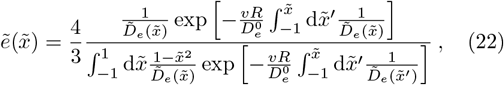

where we introduced a set of dimensionless variables, such as the dimensionless position 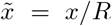, the dimensionless diffusion coefficient 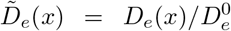, and the dimensionless enzyme profile 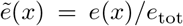 (SI Sec. IV A). We expand the enzyme profile and the velocity in the Péclet number 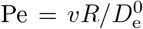. To zeroth order in the Pe, we find the adiabatic enzyme profile,

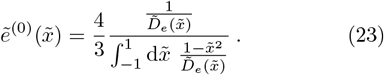

Note that 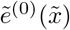 is the steady-state enzyme distribution of the enhanced diffusion equation that does not account for vesicle drift (i.e., Eq. (1)). Based on the enzyme profile 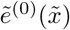, we can compute the adiabatic force,

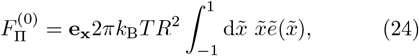

as well as the associated adiabatic velocity (via Stokes’ law),

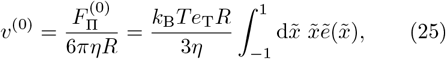

where 𝜂 denotes the viscosity of the medium in which the vesicle is submerged (SI Sec. IV B).

To linear order in Péclet number, the enzyme profile reads (SI Sec. IV A),

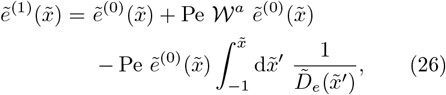

where 𝒲^*a*^ is a (Péclet-number independent) weight function that reflects the position-dependence of the effective diffusion coefficient,

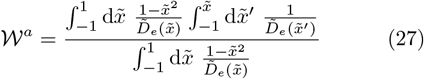

From this enzyme profile, we obtain the velocity to linear order in Péclet number,

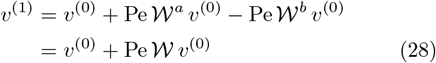

where *𝒲* = *𝒲*^*a*^−*𝒲*^*b*^, and *𝒲*^*b*^ is a weight-function similar to *𝒲*^*a*^,

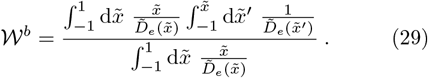

Recalling the definition of the Péclet number, we can express Eq. (28) as,

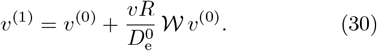

Assuming that the velocity to linear order in Péclet number, 𝑣 ^(1)^, is a good approximation of the vesicle translation velocity, 𝑣 (this is the self-consistency condition that gives the approach its name), we can solve Eq. (30) for 𝑣 = 𝑣 ^(1)^,

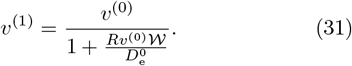

The velocity 𝑣 ^(1)^ depends on the position of the vesicle in the system, as the diffusion coefficients in the definition of *𝒲* depend on the vesicle position. Consequently, the velocity of the vesicle varies along its trajectory, and the velocity observed in the simulation is the effective vesicle velocity 𝑣 ^eff^ averaged over the positions of the vesicle along its trajectory. To compute 𝑣 ^eff^, we compute the displacement Δ*x*(*t*) = *x*(*t*) − *x*(0) of the vesicle’s position by integrating 𝑣 ^(1)^(*x*), and divide Δ*x*(*t*) the displacement by *t*,

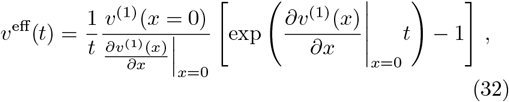

(SI Sec. IV C). This effective velocity 𝑣 ^eff^ obtained based on the velocity to linear order in Péclet number, 𝑣 ^(1)^, agrees well with the translation velocities observed in the simulations (Fig. 5). We can rationalize the dependency of the velocity on system parameters (e.g., vesicle radius, substrate gradient, basal enzyme diffusion coefficient), by studying how 𝑣 ^(1)^ depends on these parameters (SI Sec. IV D to IV F).

### Software and Data Availability

The simulation models were implemented using the JAX-MD library [61]. Simulations were run on Nvidia RTX 4090 and A100 GPUs. We simulated up to 30, 000 membrane vertices and 131, 072 enzyme particles. Additional implementation details can be found in the Supporting Information Sec. II. The code and execution instructions are available in the GitHub repository [62].

## Author Contributions

GG, HSA and UG designed the project. ESE developed the simulation code, with contributions from CLP. ESE performed the simulations, ESE and CLP analyzed the simulation data. GG, HSA and LB developed analytical theory for adiabatic translation velocities. LB extended the theory for non-adiabatic conditions via a self-consistency formalism. GG and UG supervised the project. ESE, LB, CLP, and GG wrote the manuscript, with input from all authors.

## ACKNOWLEDGMENTS

We thank Yamit Alon, Ibon Santiago and Friedrich C. Simmel for stimulating discussions. This work was supported by the Excellence Cluster ORIGINS which is funded by the Deutsche Forschungsgemeinschaft (DFG, German Research Foundation) under Germany’s Excellence Strategy - EXC-2094-390783311. Some of the simulations have been carried out on the computing facilities of the Computational Center for Particle and Astrophysics (C2PAP).

## Notes

### Competing Interest Statement

The authors have declared no competing interest.

