## Supplementary Material for "Force Generation by Enhanced Diffusion in Enzyme-Loaded Vesicles"

### Supporting Information
### Force Generation by Enhanced Diffusion in Enzyme-Loaded Vesicles

Eike S. Eberhard,* Ludwig Burger,* Cesar Lopez Pastrana,*
Hamid Seyed-Allaei, Giovanni Giunta,† and Ulrich Gerland†
*Physics of Complex Biosystems, Technical University of Munich, 85748 Garching, Germany*

#### CONTENTS

**Page**

I. Substrate Permeability through the Vesicle 2

II. Simulations 4
A. Mesh generation for the Mesh-based Vesicle Model 4
B. Numerical Integrator 5
C. Initialization of Enzyme Distribution 5
D. Analysis of Shape Parameters 5
E. Translational velocity in the mesh-based simulations 7
F. Parameters 7

III. Fluctuation Spectrum of Enzyme-filled Vesicles 8

IV. Translational Velocity of Enzyme-filled Vesicles 10
A. Steady-State Enyzme Profile 10
B. Force and Velocity 12
C. Effective Trajectory-Averaged Velocity 14
D. Substrate-Dependence of the Velocity 15
E. Radius-Dependence of the Velocity 18
F. Dependence of the Velocity on the Diffusion Coefficient 19

List of Figures 23

References 23

* These authors contributed equally to this work.
† These authors jointly supervised this work.

#### I. SUBSTRATE PERMEABILITY THROUGH THE VESICLE

In our model, we assume that the vesicle is infinitely permeable to substrate molecules. To address the question, how finite substrate permeability affects the deformation and motion of the vesicle, we investigate the relation between the internal substrate gradient and the membrane's permeability. In steady state, Fick's second law of diffusion implies that $D_s \Delta s = 0$. The solution to this Laplace equation can be obtained via an expansion in spherical harmonics. We consider the equilibrium substrate profile for a substrate gradient parallel to the $x$-axis. The boundary conditions

$$s_{\text{out}}(\mathbf{r}) = \nabla s_\infty \mathbf{e}_x \cdot \mathbf{r} = \nabla s_\infty \, x \qquad \text{for } r \gg R \tag{S1}$$
$$J = \gamma_p \left[ s_{\text{in}}(\mathbf{r}) - s_{\text{out}}(\mathbf{r}) \right] \qquad \text{for } r = R \tag{S2}$$

divide the system into two domains, the vesicle's interior and exterior. Here, $\nabla s_\infty \in \mathbb{R}$ is the external substrate gradient far away from the vesicle, and $\gamma_p$ is the permeability. The flux $J$ across the membrane is given according to Fick's first law,

$$J = -D_s \nabla s_{\text{in}} \cdot \mathbf{e}_r \Big|_{||\mathbf{r}||=R} = -D_s \nabla s_{\text{out}} \cdot \mathbf{e}_r \Big|_{||\mathbf{r}||=R} . \tag{S3}$$

We express $s_{\text{in}}(\mathbf{r})$ and $s_{\text{out}}(\mathbf{r})$ in spherical harmonics using the general ansatz for square-integrable solutions with azimuthal symmetry of the Laplace equation,

$$s(r,\theta) = \sum_{l=0}^{\infty} (A_l r^l + B_l r^{-l-1}) P_l(\cos\theta) \tag{S4}$$

where $P_l$ is the $l$-th Legendre polynomial. The coordinate system is chosen such that $x = r\cos\theta$. The terms with a negative power of $r$ inside the vesicle need to vanish since the substrate concentration is finite. Additionally, we note that the asymptotic behavior at large distances from the vesicle implies that on the outside all powers $\geq 2$ of $r$ and any constant terms must also vanish. Hence,

$$s_{\text{in}}(r,\theta) = \sum_{l=0}^{\infty} I_l r^l P_l(\cos\theta) \tag{S5}$$

$$s_{\text{out}}(r,\theta) = \nabla s_\infty r P_1(\cos\theta) + \sum_{l=0}^{\infty} O_l r^{-l-1} P_l(\cos\theta) . \tag{S6}$$

We plug our ansatz into the continuity equation (Eq. (S3)), using the identity $\nabla f \cdot \hat{e}_r \equiv \partial_r f$, and obtain

$$\sum_{l=1}^{\infty} I_l l R^{l-1} P_l = \nabla s_\infty P_1 - \sum_{l=0}^{\infty} O_l (l+1) R^{-l-2} P_l , \tag{S7}$$

which is equivalent to the conditions

$$0 = R^{-2} O_0 \tag{S8}$$
$$0 = I_1 - \nabla s_\infty + 2R^{-3} O_1 \tag{S9}$$
$$0 = l R^{l-1} I_l + (l+1) R^{-l-2} O_l \qquad \forall l \geq 2 . \tag{S10}$$

Next, we use boundary conditions (Eq. (S2)),

$$\nabla s_\infty R P_1 + \sum_{l=0}^{\infty} O_l R^{-l-1} P_l - \sum_{l=0}^{\infty} I_l R^l P_l = \frac{D_s}{\gamma_p} \sum_{l=1}^{\infty} I_l l R^{l-1} P_l, \tag{S11}$$

to derive the additional relations

$$0 = R^{-1} O_0 - I_0 , \tag{S12}$$
$$0 = \nabla s_\infty R + R^{-2} O_1 - \left( R + \frac{D_s}{\gamma_p} \right) I_1 , \tag{S13}$$
$$0 = R^{-l-1} O_l - \left( R^l + \frac{D_s}{\gamma_p} l R^{l-1} \right) I_l \qquad \forall l \geq 2 . \tag{S14}$$

We notice that both $I_0 = 0$ and $O_0 = 0$. To see that $I_l$ and $O_l$ vanish for all $l \geq 2$, we use Eq. (S14) and plug it into Eq. (S10),

$$O_l = \left(R^{2l+1} + \frac{D_s}{\gamma_p} l R^{2l}\right) I_l \qquad \forall l \geq 2 \tag{S15}$$

$$0 = \underbrace{\left(l R^{l-1} + (l+1) R^{-l-2} \left(R^{2l+1} + \frac{D_s}{\gamma_p} l R^{2l}\right)\right)}_{\geq 0} I_l \,. \tag{S16}$$

We utilize the two remaining conditions,

$$O_1 = \frac{R^3}{2} \left(\nabla s_\infty - I_1\right) \tag{S17}$$

$$0 = \nabla s_\infty R + \frac{R}{2} \left(\nabla s_\infty - I_1\right) - \left(R + \frac{D_s}{\gamma_p}\right) I_1 \tag{S18}$$

to determine

$$I_1 = \frac{\nabla s_\infty}{1 + \frac{2}{3R}\frac{D_s}{\gamma_p}} \,, \tag{S19}$$

which implies the following solution for the substrate gradient,

$$s_{\text{in}}(r,\theta) = \frac{\nabla s_\infty}{1 + \frac{2}{3R}\frac{D_s}{\gamma_p}} \, r P_1(\cos\theta) \tag{S20}$$

$$s_{\text{out}}(r,\theta) = \nabla s_\infty \, r P_1(r\cos\theta) + \nabla s_\infty \frac{R^3}{2} \frac{\frac{2D_s}{3R\gamma_p}}{1 + \frac{2D_s}{3R\gamma_p}} \frac{P_1(\cos\theta)}{r^2} \tag{S21}$$

For the internal substrate gradient to match the outside gradient, $\nabla s_{\text{in}} \approx \nabla s_{\text{out}} = \nabla s_\infty$, the permeability $\gamma_p$ must be sufficiently large,

$$\gamma_p \gg \frac{2D_s}{3R} \,. \tag{S22}$$

For urea ($D_s = 1.38\times 10^{-9}\,\text{m}^2/\text{s}$ [1, 2]) in a vesicle of radius $R = 8\,\mu\text{m}$, the right hand side evaluates to $\frac{2D_s}{3R} \approx 10^{-4}\text{m/s}$. The solubility of urea in lipids is low, implying low permeability through lipid bilayers [3]. Typical values for the permeability of urea for these artificial membranes are in the order of $\gamma_p = 10^{-8}$ m/s (Table I).

Pores can be added to the membrane to increase the permeability with regard to both water and substrate [4]. We approximate the permeability of a membrane with pores,

$$\gamma_p^s = \frac{A_{\text{pores}}^{(\text{in})}}{A_{\text{mem}}} \frac{D_s}{\delta_{\text{pore}}} + \frac{A_{\text{mem}} - A_{\text{pores}}^{(\text{out})}}{A_{\text{mem}}} \gamma_{p_0}^s \tag{S23}$$

$$= N_{\text{pores}} \frac{(r_{\text{pore}} - r_s)^2}{4R^2} \frac{D_s}{\delta_{\text{pore}}} + \left(1 - N_{\text{pores}} \frac{(R_{\text{pore}} + r_s)^2}{4R^2}\right) \gamma_{p_0}^s \,, \tag{S24}$$

where $A_{\text{pores}}^{(\text{in})}$ and $A_{\text{pores}}^{(\text{outer})}$ are the effective area of the inner channel and the area of the whole pore on the membrane, respectively. These can be approximately calculated using the inner and out pore radii $r_{\text{pore}}$ and $R_{\text{pore}}$, and the effective radius of the substrate $r_s$. The total length of the channel through which the substrate can freely diffuse is given by $\delta_{\text{pore}}$. Fig. S1a shows an illustration of the variables appearing in Eq. (S24).

In the following, we consider $\alpha$-Hemolysin as an example for pores in the membrane. It has an inner pore radius of $r_{\text{pore}} = 13\,\text{Å}$ with a total channel length of $\delta_{\text{pore}} = 100\,\text{Å}$. Its outer radius is approximately $R_{\text{pore}} = 50\,\text{Å}$ [5]. For urea we assume an effective radius of $r_s^{(\text{urea})} = 2.2\,\text{Å}$ [6]. We can use these parameters to estimate the number of pores needed to fulfill the permeability criterion (Eq. (S22)). The results are shown in Fig. S1b. We note that for $\alpha$-Hemolysin, this criterion can be satisfied if the number of pores is around $N_{\text{pores}} \approx 2 \cdot 10^5$. For this pore number, approximately 2% of the membrane area is covered by $\alpha$-Hemolysin. It is worth mentioning that other pores with a smaller area $A_{\text{pores}}^{(\text{out})}$ could be used in experiment such that the total fraction of area covered by pores would be less. Moreover, vesicles with larger radii would further relax the membrane's permeability requirements (Eq. (S22)). Giant lipid vesicles with radii of up to $R \approx 15\,\mu\text{m}$ have been realized in similar experimental settings [7].

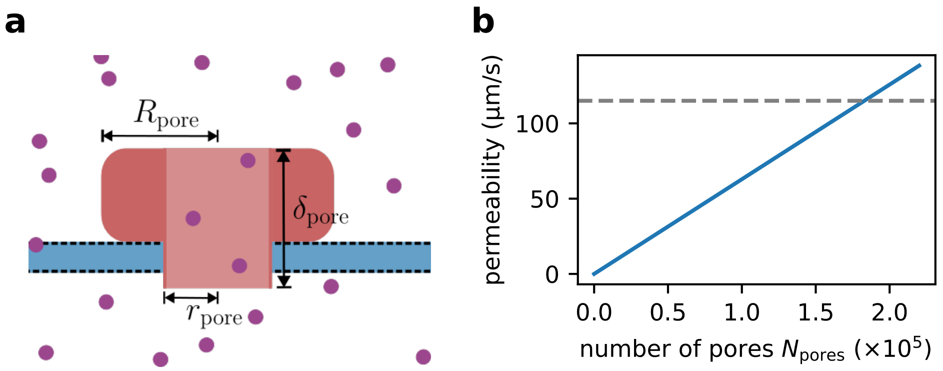

Figure S1. **Pore-dependence of membrane permeability** a) Illustration of a membrane pore. b) The membrane permeability increases linearly with the number of pores according to Eq. (S24), and exceeds the characteristic permeability (Eq. (S22), dashed line) for sufficiently high number of pores. Here, we show the permeability of urea through a POPC vesicle with a radius of $R = 8\,\mu\text{m}$ with $N_p$ $\alpha$-Hemolysin pores (Table I).

| **Lipid** | Membrane Thickness $\delta$[Å] | $\gamma_{p0}^{\text{water}}$[μm/s] | $\gamma_{p0}^{\text{urea}}$[μm/s] |
|---|---|---|---|
| **POPC** | 37.0 | 130 | 0.013 |
| **DPPC** | 28.4 | 2600 (at 50 °C) | 0.3 (at 50 °C) |
| **DOPC** | 28.8 | 158 | 0.0053 |

Table I. **Properties of Artificial Membranes (Phosphatidylcholines)** [8–14]

#### II. SIMULATIONS

##### A. Mesh generation for the Mesh-based Vesicle Model

While prominent strategies of initializing the membrane utilize the spherical geometry to formulate a structured approach in building a triangulation, this ansatz would lead to an unphysical long-ranged order in the membrane. For this reason, we choose a stochastic growth process. First, all vertices are sampled from a uniform distribution on the unit sphere. The resulting positions are then iteratively relaxed using gradient descent, w.r.t different potential energy functions. We start by applying a radial restraint term $U^{(i)}_{\text{radial restraint}} \propto (r_i - R)^6$, with $R$ the targeted equilibrium radius of the vesicle. This is combined with a repulsive soft sphere interaction potential acting between the membrane particles $U^{(\text{init})}_{\text{mem-mem}} \propto \left(1 - \frac{r}{\sigma^{(\text{init})}_{\text{mem-mem}}}\right)^4 \Theta(\sigma^{(\text{init})}_{\text{mem-mem}} - r)$, with $\sigma^{(\text{init})}_{\text{mem-mem}} \approx 2l_{\max}$. This process is run until a visually satisfying spacing between points is achieved. After this first crude relaxation step, we use the *Advancing Front Surface Reconstruction* routine based on Delaunay triangulation [15] implemented in the *Computational Geometry Algorithms Library* [16] to construct a triangulated mesh from the unstructured point cloud of vertices. This triangulation would be numerically unstable if it was combined with the simulation bond potential directly. For this reason, we further relax the triangulation with a harmonic bond potential $U^{(ij)}_{\text{harmonic-bond}} \propto (r_{ij} - l_b)^2$ and the area potential $U_{\Delta\text{-area}} = \frac{k_A}{2} \sum_{i=1}^{N_T} \frac{(A_i - A_0)^2}{A_0}$ used in the simulation (for the relaxation, the strength of the potential, $k_A$, is chosen smaller than later in the simulation). In the process of relaxation, we allow for a reconfiguration of the triangulation using the bond flipping procedure at a temperature of $T = 0$K. In the final relaxation steps, we successively add the bending potential and the volume potential and replace the harmonic bond potential with a cubic one $U^{(ij)}_{\text{cubic-bond}} \propto (r_{ij} - l_b)^3$, while increasing the area conservation coefficient $k_A$ to the correct value. After a sufficient number of steps, this results in a triangulation with very homogeneous bond lengths and triangle areas at a temperature of $T = 0\,$K, such that we can safely replace the harmonic potential with the correct bond potential. This completes the membrane potential in the simulation. The resulting membrane state is then equilibrated to a temperature of $T = 300$K. For the simulation, the cubic bond potential is replaced by the bond potential introduced in Eq. (14).

##### B. Numerical Integrator

In the dilute regime, enzyme-enzyme interactions are negligible, and the concentration profile $e$ in the bulk is proportional to the single-particle probability distribution $e(\mathbf{r},t) \propto P(\mathbf{r},t)$. We use the theorem

$$\partial_t P(\mathbf{r},t) = -\nabla_{\mathbf{r}} \left[A(\mathbf{r})P\right] + \frac{1}{2}\nabla^2_{\mathbf{r}} \left[(C(\mathbf{r}))^2 P\right] \overset{\text{Itô}}{\Longleftrightarrow} \frac{d\mathbf{r}}{dt} = A(\mathbf{r}) + C(\mathbf{r})\eta(t) \,, \tag{S25}$$

to express the Fokker-Planck equation for the probability distribution as a stochastic differential equation (Langevin equation) assuming a zero-mean unit-variance white noise process $\eta(t)$ [17, 18]. The theorem holds for Itô's interpretation of stochastic calculus, which is applicable to chemical reactions. For $A(\mathbf{r}) = 0$ and $C(\mathbf{r}) = \sqrt{2D_{\mathrm{e}}(\mathbf{r})}$, we obtain the Brownian dynamics equation

$$\frac{d\mathbf{r}}{dt} = \sqrt{2D_{\mathrm{e}}(\mathbf{r})}\eta(t) \,. \tag{S26}$$

To account for mechanistic interactions of the enzymes with the vesicle boundary, we need to introduce an additional force term

$$\frac{d\mathbf{r}_e}{dt} = \frac{1}{\gamma_e}\mathbf{F} + \sqrt{2D_{\mathrm{e}}(\mathbf{r})}\eta(t) \,. \tag{S27}$$

For the purpose of this work we make the assumption that Einstein's relation is fulfilled. Hence, the friction reads

$$\gamma_e(\mathbf{r}) = \frac{k_B T}{D_{\mathrm{e}}(\mathbf{r})} \,, \tag{S28}$$

and we arrive at the Brownian dynamics equation

$$\frac{d\mathbf{r}_e}{dt} = \frac{D_{\mathrm{e}}(\mathbf{r})}{k_B T}\mathbf{F} + \sqrt{2D_{\mathrm{e}}(\mathbf{r})}\eta(t) \tag{S29}$$

which we use for the simulation. This governing equation is compatible with enhanced diffusion that emerges due to a passive mechanism, e.g., a change in the hydrodynamic radius of the enzyme induced by substrate binding [19]. To model enhanced diffusion due to active enzyme leaps, one would need to use a modified governing equation [20], in which the friction coefficient, $\gamma = \frac{k_{\mathrm{B}}T}{D^0_{\mathrm{e}}}$, is independent of substrate concentration,

$$\frac{d\mathbf{r}_e}{dt} = \frac{D^0_{\mathrm{e}}}{k_B T}\mathbf{F} + \sqrt{2D_{\mathrm{e}}(\mathbf{r})}\eta(t) \,. \tag{S30}$$

In our model no forces are acting on the enzymes in the vesicle. Therefore, both microscopic pictures are expected to yield the same concentration profile.

##### C. Initialization of Enzyme Distribution

In the mesh-based simulation, the enzymes are initialized following the adiabatic steady-state distribution. The enzymes are placed within a slightly smaller $R$ than the vesicle to avoid numerical instabilities associated with the repulsion force between enzymes and the vertices of the mesh.
In the simplified simulation, the positions of the enzymes are initialized randomly, resulting in a homogeneous distribution within the volume of the vesicle. Control runs with adiabatic steady-state initialization revealed no dependence of the results on the initial distribution of the enzymes when allowing for sufficient equilibration times, typically in the order of $\sim 200\,\mathrm{ms}$.

##### D. Analysis of Shape Parameters

To characterize the shape of the vesicle (and the change in shape due to enhanced diffusion), we resort to a set of order parameters defined in terms of the gyration tensor,

$$Q_{\alpha\beta} = \frac{1}{N}\sum_{i=1}^{N_V}(\mathbf{r}^{(i)}_\alpha - \mathbf{R}^{(i)}_\alpha)(\mathbf{r}^{(i)}_\beta - \mathbf{R}^{(i)}_\beta), \tag{S31}$$

where $\alpha$ and $\beta$ denote the components $x, y, z$ of position $\mathbf{r}^i$ of the $i$th mesh vertex. The sum runs over all the $N_V$ vertices. The term $\mathbf{R}$ is the center of mass of the vesicle. Via diagonalization, the tensor $\mathbf{Q}$ is uniquely defined by its eigenvectors $\hat{\mathbf{e}}_1$, $\hat{\mathbf{e}}_2$, $\hat{\mathbf{e}}_3$ and the associated eigenvalues $\lambda_1 \leq \lambda_2 \leq \lambda_3$.
The asphericity $\mathcal{A}$ quantifies the degree of non-spherical distribution of the mass. It is defined as [21, 22, 41]

$$\mathcal{A} \equiv \frac{(\lambda_1 - \lambda_2)(\lambda_2 - \lambda_3)(\lambda_3 - \lambda_1)}{2(\lambda_1 + \lambda_2 + \lambda_3)^2}. \tag{S32}$$

The asphericity is normalized such that $0 \leq \mathcal{A} \leq 1$, where $\mathcal{A} = 0$ zero indicates a perfect spherical symmetry (not necessarily spherical in shape), and $\mathcal{A} = 1$ a one-dimensional distribution of mass (e.g., a cylinder).
The prolateness $\mathcal{P}$ allows to discriminate spheroids with prolate or oblate morphology [41],

$$\mathcal{P} \equiv \frac{(2\lambda_1 - \lambda_2 - \lambda_3)(2\lambda_2 - \lambda_1 - \lambda_3)(2\lambda_3 - \lambda_1 - \lambda_2)}{2(\lambda_1^2 + \lambda_2^2 + \lambda_3^2 - \lambda_1\lambda_2 - \lambda_1\lambda_3 - \lambda_2\lambda_3)^{3/2}} \tag{S33}$$

The prolateness is defined in the range $-1 \leq \mathcal{P} \leq 1$, where $\mathcal{P} < 0$ for oblate shapes and $\mathcal{P} > 0$ for prolate spheroids. We define the prolateness axis $\mathcal{P}_x = |\hat{\mathbf{e}}_1 \cdot \hat{\mathbf{e}}_x|$, where $\hat{\mathbf{e}}_1$ is the eigenvector associated to the eigenvalue $\lambda_1$ and $\hat{\mathbf{e}}_x$ is the orientation vector of the substrate gradient.
The observables described above rely on access to the 3D configuration of the vesicle. However, most experimental observations of vesicles use light microscopy imaging and can only capture the 2D cross-sections along the equatorial plane. We characterize the shape of the cross-section via its ellipticity,

$$\varepsilon = \frac{a}{b} - 1. \tag{S34}$$

Here, $a$ denotes the length of the ellipse along the direction of the substrate gradient, while $b$ denotes the length of the axis perpendicular to the direction of the gradient. We pick the direction along which $b$ is measured such that $b$ is maximal. A comparison of the shape parameters for vesicles under hyper- and hypoosmotic conditions is shown in Fig. S2.

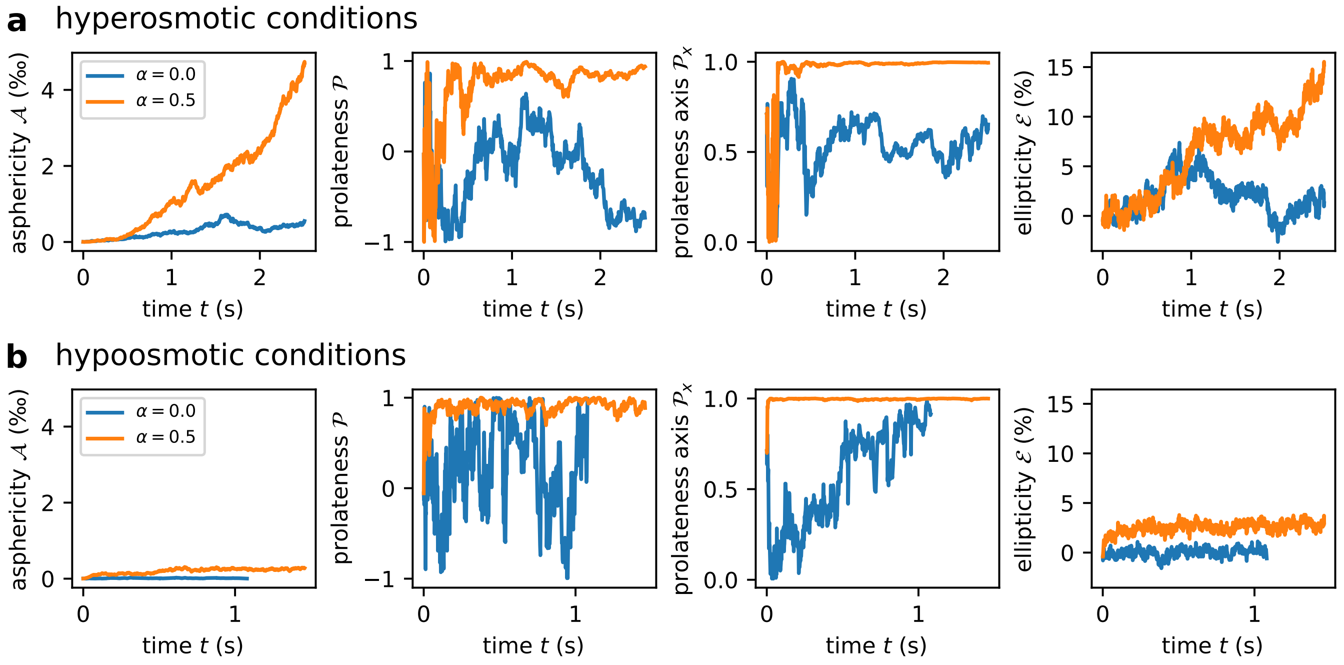

Figure S2. **Shape parameters of vesicles in hyper- and hypoosmotic conditions**. Asphericity $\mathcal{A}$, prolateness $\mathcal{P}$, prolateness axis $\mathcal{P}_x$ and ellipticity $\mathcal{E}$ for vesicles under hyperosmotic (a) and hypoosmotic conditions (b). In the case of hyperosmotic conditions, low surface tension is achieved by reducing the desired volume below the native sphere volume, $\hat{V} = V/V_0 = 0.9$ with volume stiffness $k_V = 129\,\mathrm{N\,m^{-2}}$ (Method section). Under hypoosmotic conditions, no volume constraint is imposed, $k_V = 0\,\mathrm{N\,m^{-2}}$. While asphericity and ellipticity saturate in the case of hypoosmotic conditions, they continue to increase throught the entire simulated timeframe. The parameters used in the simulations are summarized in Table II.

##### E. Translational velocity in the mesh-based simulations

Simulating the time-evolution of the mesh-based vesicle model (Methods Section) for the parameters presented in Table II (parameters chosen based on [7]) reveals that the translation velocity observed in the mesh-based simulation is smaller than the velocity expected based the force **F** observed in the simulation if we assume that the friction is set by Stokes' law for a vesicle of radius $R$ in a medium with viscosity similar to that of water (Fig. S3). The problem is caused by the choice of the diffusion coefficient of the membrane vertices, $D_{\mathrm{mem}}$, as these coefficients model the viscosity of the medium implicitly. The drag coefficient $\gamma_{\mathrm{mem}}$ for a membrane vertex is set by the vertex diffusion coefficient via the Einstein relation,

$$\gamma_{\mathrm{mem}} = \frac{k_{\mathrm{B}}T}{D_{\mathrm{mem}}}. \tag{S35}$$

Thus, the friction force acting on a single vertex equals $f_i = \gamma_{\mathrm{mem}}\, v = (k_{\mathrm{B}}T)\,/\,(D_{\mathrm{mem}})v$ (for vesicle velocity $v$), and the total friction acting on all vertices amounts to

$$F_{\mathrm{drag,\,vertex}} = \sum_{i=1}^{N_V} f_i = \frac{k_{\mathrm{B}}T}{D_{\mathrm{mem}}} N_V v \tag{S36}$$

Enforcing that this drag force is equal to the drag force expected based on Stokes' law,

$$F_{\mathrm{drag\,Stokes}} = 6\pi\eta R v, \tag{S37}$$

implies that the membrane vertex diffusion coefficient needs to be chosen as

$$D_{\mathrm{mem}} = \frac{k_B T N_V}{6\pi\eta R}. \tag{S38}$$

The value $D_{\mathrm{mem}}$ used in the mesh-based simulation is $\approx 37$ times smaller than the desired value according to Eq. (S38) (see Table II and [7]). However, choosing smaller $D_{\mathrm{mem}}$ is challenging due to numerical stability: Using the correct value for $D_{\mathrm{mem}}$ requires reducing the integration time step. Albeit this should lead to the response expected for a vesicle in water, the reduction of $dt$ makes the execution of the simulation of a duration in the order of months to years. This long time is a requirement for the equilibration of the enzyme distribution, and therefore this approach is computationally not feasible.
We validated the notion introduced in Eq. (S38) using the simplified simulation: Following Eq. (S38), we set the viscosity in the simplified vesicle model such that it matches the vertex diffusion coefficient used in the full simulation (Table II). With this choice of viscosity, we find that the translation velocity observed in mesh-based and simplified simulation agree well (Fig. S3).

##### F. Parameters

| Parameters | Model Units | Physical units |
|---|---|---|
| principal properties | | |
| vesicle radius in equilibrium $R$ | 32 | 8.0 µm |
| thermal energy unit $k_{\mathrm{B}}T$ | 0.2 | $4.14 \times 10^{-21}$ J |
| time scale $\tau$ | $1.25 \times 10^5$ | 7.3 s |
| Michaelis-Menten-constant $K_M$ | $2.8 \times 10^4$ | 3.0 mM |
| Vesicle properties | | |
| number of vertices $N_V$ | 30000 | 30000 |
| number of triangles $N_T$ | $2(N_V - 2)$ | $2(N_V - 2)$ |
| bending rigidity $\kappa$ | $20\, k_{\mathrm{B}}T$ | $8.28 \times 10^{-20}$ J |
| average bond length $l_b$ | $4R\sqrt{\pi/(N_T\sqrt{3})}$ | 0.176 µm |
| repulsive bond stiffness $k_B^{\mathrm{rep}}$ | $2046\, k_{\mathrm{B}}T$ | $8.47 \times 10^{-18}$ J/m |
| attractive bond stiffness $k_B^{\mathrm{att}}$ | $10\, k_B^{\mathrm{rep}}$ | $8.47 \times 10^{-17}$ J/m |
| repulsive stiffness $\lambda^{\mathrm{rep}}$ | 0.53 | 0.53 |
| attractive stiffness $\lambda^{\mathrm{att}}$ | 0.28 | 0.28 |
| minimum bond length $l_{\mathrm{min}}$ | $0.6\, l_b$ | 0.11 µm |
| potential cutoff length $l_{c_1}$ | $0.8\, l_b$ | 0.14 µm |

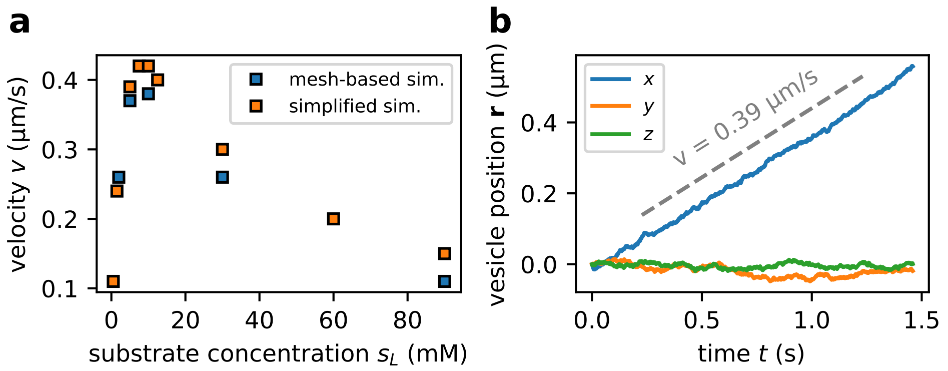

Figure S3. **Translation of enyzme-loaded vesicles** a) The translation velocities observed in the simplified simulation (blue) and full mesh-based simulation (orange) agree well, provided the viscosity of the medium in the simplified simulation is chosen to match the vertex diffusion coefficient of the mesh-based model. b) Time-evolution of the vesicle position (center of mass) in the full mesh-based simulation. The vesicle moves downstream along the substrate gradient with velocity $v \approx 0.39\,\mu\mathrm{m\,s}^{-1}$ (determined via linear fit). The vesicle is in its high surface-tension state (i.e., no volume constraint). The parameters used for the simulation are listed in Table II. Note that the translation velocity is lower than the one observed in the simplified simulation (Fig. 4). The parameters for the full mesh-based simulation are summarized in Table II (same parameters as used in [7]), and the viscosity is computed via Eq. (S38). Note that this viscosity $\eta_{\mathrm{used}} \approx 37\,\mathrm{mPa\,s}$ is about 40 times larger than the true viscosity of water, $\eta_{\mathrm{water}} = 1\,\mathrm{mPa\,s}$.

| | | |
|---|---|---|
| potential cutoff length $l_{c_0}$ | $1.2\,l_b$ | $0.21\,\mu\mathrm{m}$ |
| maximum bond length $l_{\max}$ | $1.4\,l_b$ | $0.25\,\mu\mathrm{m}$ |
| desired vesicle area $A$ | $4\pi R^2$ | $8.04 \times 10^2\,\mu\mathrm{m}^2$ |
| desired vesicle volume $V_0$ | $4\pi R^3/3$ | $2.14 \times 10^3\,\mu\mathrm{m}^3$ |
| local area stiffness $k_A$ | $6.43 \times 10^6\,k_{\mathrm{B}}T/A$ | $3.3 \times 10^{-5}\,\mathrm{J/m}^2$ |
| volume stiffness $k_V$ (hyperosmotic condition) | $1.6 \times 10^7\,k_{\mathrm{B}}T/R^3$ | $129\,\mathrm{J/m}^3$ |
| target volume $\hat{V}$ (hyperosmotic condition) | $0.9V_0$ | $1.93 \times 10^3\,\mu\mathrm{m}^3$ |
| volume stiffness $k_V$ (hypoosmotic condition) | $0\,k_{\mathrm{B}}T/R^3$ | $0\,\mathrm{J/m}^3$ |
| membrane diffusion coefficient $D_{\mathrm{m}}$ | $2.5\,R^2/\tau$ | $21.9\,\mu\mathrm{m}^2\,\mathrm{s}^{-1}$ |
| flipping frequency $\omega$ | $6.4 \times 10^6/\tau$ | $8.8 \times 10^5\,\mathrm{s}^{-1}$ |
| flipping probability $\psi$ | 0.3 | 0.3 |
| enzyme properties | | |
| number of enzymes $N_e$ | $131,072$ | $101.5\,\mathrm{nM} \sim 100\,\mathrm{nM}$ |
| number of enzymes $N_e$ in Fig. 2 and S2 under hyperosmotic conditions | $55,296$ | $50.5\,\mathrm{nM} \sim 50\,\mathrm{nM}$ |
| equilibrium diffusion coefficient $D_0$ | $3.6 \times 10^{-2}$ | $39\,\mu\mathrm{m}^2\,\mathrm{s}^{-1}$ |
| enhanced diffusion factor $\alpha$ | 0.5 | 0.5 |
| enz-mem interaction radius $\sigma_{\mathrm{enz}}$ | $1.5\,l_b$ | $0.26\,\mu\mathrm{m}$ |
| enz-mem interaction energy param. $\epsilon$ | $40\,k_{\mathrm{B}}T$ | $1.66 \times 10^{-19}\,\mathrm{J}$ |
| environmental properties | | |
| maximum substrate concentration $s_{\max}$ | $3.33\,K_M$ | $10.0\,\mathrm{mM}$ |
| system length $L$ | $2.4\,R$ | $19.2\,\mu\mathrm{m}$ |
| further parameters | | |
| time step size $dt$ | $2.5 \times 10^{-5}$ | $1.46\,\mathrm{ns}$ |
| total simulation time | $2.5 \times 10^4$ | $1.46\,\mathrm{s}$ |

Table II: Default parameter set used in mesh-based simulations. Principal properties and vesicle properties are the same as in [7] except for a subset of the parameters associated with the bond potential. In individual runs, some of these parameters were varied. In these cases, the affected parameters are stated explicitly, and the remaining independent parameters remain unchanged. The equilibrium enzyme diffusion coefficient and the Michaelis-Menten constant $K_{\mathrm{M}}$ are chosen based on the values measured for urease by Jee et al. [23].

#### III. FLUCTUATION SPECTRUM OF ENZYME-FILLED VESICLES

We aim to compare the fluctuation spectrum observed in the simulation (without enhanced diffusion) to the fluctuation spectrum predicted by theory [7]. To this end, we need to determine the surface tension of vesicle based on the membrane parameters used in the model. Excluding the volume potential, the membrane energy in our system is

| Parameters | Model Units | Physical units |
|---|---|---|
| principal properties | | |
| vesicle radius $R$ | 32 | 8.0 µm |
| thermal energy unit $k_{\mathrm{B}}T$ | 0.2 | $4.14 \times 10^{-21}$ J |
| Michaelis-Menten-constant $K_M$ | $2.8 \times 10^4$ | 3.0 mM |
| vesicle properties | | |
| vesicle friction $\gamma_{\mathrm{ves}}$ | $6\pi\eta R$ | $1.5 \times 10^{-7}$ Ns/m |
| vesicle diffusion coefficient $D_{\mathrm{ves}}$ | $\frac{k_{\mathrm{B}}T}{\gamma_{\mathrm{ves}}}$ | $2.8 \times 10^{-2}$ µm$^2$ s$^{-1}$ |
| shell stiffness $k_{\mathrm{ves}}$ | 50 $k_{\mathrm{B}}T$ | $3.3 \times 10^{-6}$ J/m$^2$ |
| shell stiffness $k_{\mathrm{ves}}$ in Fig. S3 | 5 $k_{\mathrm{B}}T$ | $3.3 \times 10^{-7}$ J/m$^2$ |
| shell stiffness $k_{\mathrm{ves}}$ for static vesicle in Fig. S4 | 5 $k_{\mathrm{B}}T$ | $3.3 \times 10^{-7}$ J/m$^2$ |
| enzyme properties | | |
| number of enzymes $N_e$ | 131072 | 101.5 nM $\sim$ 100 nM |
| equilibrium diffusion coefficient $D_0$ | $3.6 \times 10^{-2}$ | 39 µm$^2$ s$^{-1}$ |
| enhanced diffusion factor $\alpha$ | 0.5 | 0.5 |
| environmental properties | | |
| maximum substrate concentration $s_{\mathrm{max}}$ | 3.33 $K_M$ | 10 mM |
| maximum substrate concentration $s_{\mathrm{max}}$ in Fig. S4 | 1.66 $K_M$ | 5 mM |
| system length $L$ | 76.8 | 19.2 µm |
| viscosity of water $\eta$ | 12.9 | $1.0 \times 10^{-3}$ Pa s |
| effective viscosity $\eta$ in Fig. S3 | 477.3 | $37 \times 10^{-3}$ Pa s |
| high viscosity $\eta$ in Figs. S5, S7, S9, S11 | 12900 | 1.0 Pa s |
| general parameters | | |
| time step size $dt$ | 0.0051 | 300 ns |
| total simulation time | $2.5 \times 10^4$ | 1.5 s |

Table III. Default parameter set used in simplified simulations. The equilibrium diffusion coefficient and the Michaelis-Menten constant $K_M$ are the values measured for urease by Jee et al. [23].

given by

$$E_m = E_b + E_s, \tag{S39}$$

where $E_b$ is the bending term and $E_s$ is the area stretching term. The bending potential is given by part of the Helfrich Hamiltonian,

$$E_b = 2\kappa \iint H^2 dS\,, \tag{S40}$$

where $\kappa$ is the bending stiffness and $H$ is the mean curvature. For spherical vesicles, $E_b = 8\pi\kappa$. The area stretching energy equals

$$E_s = \frac{k_s}{2} \sum_{t=1}^{N_t} \frac{(A_t - A_0)^2}{A_0}, \tag{S41}$$

where $N_t$ is the total number of triangles of the mesh, $A_t$ is the current triangle area and $A_0$ is the characteristic rest area of the triangles. If we consider a spherical shell discretized with triangles of equal area, the expression can be simplified to

$$E_s = 2\pi k_s \frac{\left(R^2 - R_0^2\right)^2}{R_0^2}. \tag{S42}$$

This expression is equivalent to the expected stretching energy cost of a continuum material.

The force due to the membrane potential in the absence of enzymes equals $f_m = f_s + f_b$. We find the force along the radial direction by computing the radial component of the gradient of $E_m$,

$$f_m = -\partial_R E_m = \frac{8\pi R k_s \left(R_0^2 - R^2\right)}{R_0^2}. \tag{S43}$$

It is worth noting that the bending term does not contribute to the total force on the radial direction since $\partial_R E_b = 0$. In addition to the forces due to the membrane potential, we also need to account for the force exerted by enzymes. The osmotic pressure associated with the enzymes is given by the ideal gas law, $\Pi = k_B T \left(\frac{N_e}{V}\right)$, where $N_e$ is the number of enzymes. Since the pressure equals force per area, $\Pi = f/S$, we can determine the force exerted by the enzymes in a spherical vesicle ($S = 4\pi R^2$, $V = 4/3\pi R^3$),

$$f_{\text{enz}} = \frac{3n_e k_B T}{R}\,. \tag{S44}$$

In equilibrium, the forces are balanced, i.e., the force due to the membrane potential and the force exerted by the enzymes need to add up to zero, $f_m(R^*) + f_{\text{enz}}(R^*) = 0$. This condition sets the equilibrium vesicle radius,

$$R^* = R_0 \sqrt{\frac{1 + \sqrt{1 + \frac{3N_e k_\text{B} T}{2\pi R_0^2 k_s}}}{2}}\,. \tag{S45}$$

To determine the surface tension $\sigma$, we resort to the Young-Laplace equation. For an isotropically pressurized spherical shell, the pressure in the shell is related to the surface tension via $\Pi = 2\sigma/R$. Recalling that the only internal force of the membrane is associated with the area stretching (since $f_b = 0$) allows us to express the pressure as a function of $f_s$, $\Pi = f_s/(4\pi R^2)$. Consequently, the surface tension equals,

$$\sigma = \left.\frac{f_s(R)}{8\pi R}\right|_{R=R^*} = k_s\left(1 - \frac{(R^*)^2}{R_0^2}\right) = \frac{k_s - k_s\sqrt{1 + \frac{3N_e k_\text{B} T}{2\pi R_0^2 k_s}}}{2} \approx -0.97\,\mu\text{N}\,\text{m}^{-1}\,. \tag{S46}$$

Having determined the surface tension $\sigma$ and the equilibrium radius $R^*$, we can compute the resulting fluctuation spectrum for the bending stiffness used in the simulation ($\kappa = 20\ k_\text{B}T$) [7],

$$\left\langle |a_\ell|^2 \right\rangle = \frac{k_B T}{4\pi\sigma}\left(\frac{1}{\ell} - \frac{1}{\sqrt{\frac{\sigma}{\kappa}R^2 + \ell^2}}\right), \tag{S47}$$

where the average radius is $R = R^*$. We find very good agreement between the fluctuation spectrum obtained in simulations and that from our theoretical predictions (Fig. 3).

#### IV. TRANSLATIONAL VELOCITY OF ENZYME-FILLED VESICLES

In order to compute the translational velocity of an enzyme-filled vesicle, we need to compute (i) the steady-state concentration profile of the enzymes in the vesicle, as well as (ii) the resulting force acting on the vesicle and the associated velocity. However, the steady-state enzyme profile depends on the translation velocity of the vesicle, implying that it is necessary to know the velocity of the vesicle in order to compute the velocity via the steady-state enzyme profile. We use this notion to formulate a self-consistency condtion for the velocity of the vesicle. Together with an expansion in small Péclet number, the self-consistency approach allows us to derive a closed expression for the vesicle velocity.

##### A. Steady-State Enyzme Profile

For the derivation of the steady-state enzyme profile, we start from the enhanced diffusion equation that accounts for the motion of the enzyme (Eq. (4)). We assume that the enzyme concentration only depends on the position along the direction of the substrate, which simplifies Eq. (4) in steady-state to

$$\partial_t e = 0 = \partial_x^2(D_e(x)e(x)) + v\,\partial_x e(x). \tag{S48}$$

The solution of this differential equation reads

$$e(x) = \frac{e(-R)D_e(-R)}{D_e(x)} \exp\left[-v\int_{-R}^{x} \mathrm{d}x' \frac{1}{D_e(x')}\right]. \tag{S49}$$

As we aim to express the enzyme profile as a function of the total concentration, we need to relate $e(-R)$ to the total number of enzymes,

$$N_{\mathrm{T}} = \int \mathrm{d}V e_{\mathrm{T}} = \frac{4\pi R^3}{3} e_{\mathrm{T}} \tag{S50}$$

$$N_{\mathrm{T}} = \int \mathrm{d}V e(x) = e(-R) D_e(-R) \int_{-R}^{R} \mathrm{d}x \frac{\pi(R^2 - x^2)}{D_e(x)} \exp\left[-v \int_{-R}^{x} \mathrm{d}x' \frac{1}{D_e(x')}\right] \tag{S51}$$

To simplify further, we rescale $x$ in units of radius, i.e., $\tilde{x} = \frac{x}{R}$,

$$N_{\mathrm{T}} = e(-R) D_e(-R) \pi R^3 \int_{-1}^{1} \mathrm{d}\tilde{x} \frac{1 - \tilde{x}^2}{D_e(\tilde{x})} \exp\left[-v \int_{-1}^{\tilde{x}} \mathrm{d}\tilde{x}' \frac{R}{D_e(\tilde{x}')}\right] \tag{S52}$$

Solving for $e(-R)$, we find

$$e(-R) = \frac{\frac{4}{3}\pi R^3 e_{\mathrm{T}}}{D_e(-R)\pi R^3 \int_{-1}^{1} \mathrm{d}\tilde{x} \frac{1-x^2}{D_e(\tilde{x})} \exp\left[-v \int_{-1}^{\tilde{x}} \mathrm{d}\tilde{x}' \frac{R}{D_e(\tilde{x}')}\right]} \tag{S53}$$

$$= \frac{4 e_{\mathrm{T}}}{3 D_e(-R) \int_{-1}^{1} \mathrm{d}\tilde{x} \frac{1-x^2}{D_e(\tilde{x})} \exp\left[-v \int_{-1}^{\tilde{x}} \mathrm{d}\tilde{x}' \frac{R}{D_e(\tilde{x}')}\right]} \tag{S54}$$

We can plug this back into the formula for the enzyme profile $e(x)$,

$$e(x) = \frac{4 e_{\mathrm{T}} \frac{1}{D_e(x)} \exp\left[-v \int_{-R}^{x} \mathrm{d}x' \frac{1}{D_e(x')}\right]}{3 \int_{-1}^{1} \mathrm{d}\tilde{x} \frac{1-x^2}{D_e(\tilde{x})} \exp\left[-v \int_{-1}^{\tilde{x}} \mathrm{d}\tilde{x}' \frac{R}{D_e(\tilde{x}')}\right]}. \tag{S55}$$

For consistency, we also rescale the position $x$ to the dimensionless position $\tilde{x}$ in the numerator,

$$e(\tilde{x}) = \frac{4}{3} e_{\mathrm{T}} \frac{\frac{1}{D_e(\tilde{x})} \exp\left[-v \int_{-1}^{\tilde{x}} \mathrm{d}\tilde{x}' \frac{1}{D_e(\tilde{x}')}\right]}{\int_{-1}^{1} \mathrm{d}\tilde{x} \frac{1-\tilde{x}^2}{D_e(\tilde{x})} \exp\left[-v \int_{-1}^{\tilde{x}} \mathrm{d}\tilde{x}' \frac{R}{D_e(\tilde{x}')}\right]}. \tag{S56}$$

To make the representation of the enzyme profile fully dimensionless, we introduce the dimensionless enzyme concentration, $\tilde{e}(\tilde{x}) = e(\tilde{x})/e_{\mathrm{T}}$, as well as the dimensionless diffusion coefficient, $\tilde{D}_e(\tilde{x}) = D_e(\tilde{x})/D_{\mathrm{e}}^0$,

$$\tilde{e}(\tilde{x}) = \frac{4}{3} \frac{\frac{1}{\tilde{D}_e(\tilde{x})} \exp\left[-\frac{vR}{D_{\mathrm{e}}^0} \int_{-1}^{\tilde{x}} \mathrm{d}\tilde{x}' \frac{1}{\tilde{D}_e(\tilde{x})}\right]}{\int_{-1}^{1} \mathrm{d}\tilde{x} \frac{1-\tilde{x}^2}{\tilde{D}_e(\tilde{x})} \exp\left[-\frac{vR}{D_{\mathrm{e}}^0} \int_{-1}^{\tilde{x}} \mathrm{d}\tilde{x}' \frac{1}{\tilde{D}_e(\tilde{x}')}\right]} \tag{S57}$$

The dimensionless representation allows us to identify the Péclet number, $\mathrm{Pe} = vR/D_{\mathrm{e}}^0$. Even for the highest velocities observed in the simulation ($v \approx 0.6\,\mathrm{\mu m\,s^{-1}}$ for a vesicle with radius $R = 8\,\mathrm{\mu m}$ and diffusion coefficient $D_{\mathrm{e}}^0 = 39\,\mathrm{\mu m^2\,s^{-1}}$), the Péclet number is small, $\mathrm{Pe} \approx 0.12$, which allows us to expand the enzyme profile, $\tilde{e}(\tilde{x})$ in the Péclet number.

To zeroth order, i.e., taking the limit $\mathrm{Pe} \to 0$, we find the adiabatic enzyme profile,

$$\tilde{e}^{(0)}(\tilde{x}) = \frac{4}{3} \frac{\frac{1}{\tilde{D}_e(\tilde{x})}}{\int_{-1}^{1} \mathrm{d}\tilde{x} \frac{1-\tilde{x}^2}{\tilde{D}_e(\tilde{x})}}. \tag{S58}$$

Note that this profile is the solution to the enhanced diffusion equation that does not account for the vesicle drift (Eq. (1)).

To obtain the enzyme profile to linear order in Pe, we expand the exponential functions in small arguments,

$$\tilde{e}(\tilde{x}) \approx \frac{4}{3} \frac{\frac{1}{\tilde{D}_e(\tilde{x})}\left[1-\mathrm{Pe}\int_{-1}^{\tilde{x}} \frac{1}{\tilde{D}_e(\tilde{x}')}\mathrm{d}\tilde{x}'\right]}{\int_{-1}^{1}\mathrm{d}\tilde{x}\;\frac{1-\tilde{x}^2}{D_e(\tilde{x})}\left[1-\mathrm{Pe}\int_{-1}^{\tilde{x}}\mathrm{d}\tilde{x}'\frac{1}{\tilde{D}_e(\tilde{x})}\right]} \tag{S59}$$

$$= \frac{4}{3} \frac{\frac{1}{\tilde{D}_e(\tilde{x})}}{\int_{-1}^{1}\mathrm{d}\tilde{x}\;\frac{1-\tilde{x}^2}{D_e(\tilde{x})}\left[1-\mathrm{Pe}\int_{-1}^{\tilde{x}}\mathrm{d}\tilde{x}'\frac{1}{\tilde{D}_e(\tilde{x})}\right]} - \frac{4}{3}\mathrm{Pe}\frac{\frac{1}{\tilde{D}_e(\tilde{x})}\int_{-1}^{\tilde{x}}\mathrm{d}\tilde{x}'\frac{1}{\tilde{D}_e(\tilde{x}')}}{\int_{-1}^{1}\mathrm{d}\tilde{x}\;\frac{1-\tilde{x}^2}{D_e(\tilde{x})}\left[1-\mathrm{Pe}\int_{-1}^{\tilde{x}}\mathrm{d}\tilde{x}'\frac{1}{\tilde{D}_e(\tilde{x})}\right]} \tag{S60}$$

$$= \frac{4}{3}\frac{\frac{1}{\tilde{D}_e(\tilde{x})}}{\int_{-1}^{1}\mathrm{d}\tilde{x}\frac{1-\tilde{x}^2}{\tilde{D}_e(\tilde{x})}} + \frac{4}{3}\frac{\frac{1}{\tilde{D}_e(\tilde{x})}}{\int_{-1}^{1}\mathrm{d}\tilde{x}\;\frac{1-\tilde{x}^2}{\tilde{D}_e(\tilde{x})}}\mathrm{Pe}\frac{\int_{-1}^{1}\mathrm{d}\tilde{x}\;\frac{1-\tilde{x}^2}{\tilde{D}_e(\tilde{x})}\int_{-1}^{\tilde{x}}\mathrm{d}\tilde{x}'\frac{1}{\tilde{D}_e(\tilde{x}')}}{\int_{-1}^{1}\mathrm{d}\tilde{x}\frac{1-\tilde{x}^2}{\tilde{D}_e(\tilde{x})}} \tag{S61}$$

$$-\frac{4}{3}\frac{\frac{1}{\tilde{D}_e(\tilde{x})}}{\int_{-1}^{1}\mathrm{d}\tilde{x}\;\frac{1-\tilde{x}^2}{\tilde{D}_e(\tilde{x})}}\mathrm{Pe}\int_{-1}^{\tilde{x}}\mathrm{d}\tilde{x}'\frac{1}{\tilde{D}_e(\tilde{x}')} + \mathcal{O}\left((\mathrm{Pe})^2\right) \tag{S62}$$

$$= \tilde{e}^{(0)}(\tilde{x}) + \mathrm{Pe}\;\tilde{e}^{(0)}(\tilde{x})\;\mathcal{W}^a - \mathrm{Pe}\;\tilde{e}^{(0)}(\tilde{x})\int_{-1}^{\tilde{x}}\mathrm{d}\tilde{x}'\;\frac{1}{\tilde{D}_e(\tilde{x})} + \mathcal{O}\left((\mathrm{Pe})^2\right), \tag{S63}$$

where we introduced a Péclet-number independent weight $\mathcal{W}^a$,

$$\mathcal{W}^a = \frac{\int_{-1}^{1}\mathrm{d}\tilde{x}\;\frac{1-\tilde{x}^2}{\tilde{D}_e(\tilde{x})}\int_{-1}^{\tilde{x}}\mathrm{d}\tilde{x}'\frac{1}{\tilde{D}_e(\tilde{x}')}}{\int_{-1}^{1}\mathrm{d}\tilde{x}\frac{1-\tilde{x}^2}{\tilde{D}_e(\tilde{x})}}. \tag{S64}$$

Thus, the enzyme profile to linear order in Péclet number reads

$$\tilde{e}^{(1)}(\tilde{x}) = \tilde{e}^{(0)}(\tilde{x}) + \mathrm{Pe}\;\tilde{e}^{(0)}(\tilde{x})\;\mathcal{W}^a - \mathrm{Pe}\;\tilde{e}^{(0)}(\tilde{x})\;\int_{-1}^{\tilde{x}}\mathrm{d}\tilde{x}'\;\frac{1}{\tilde{D}_e(\tilde{x})}\,. \tag{S65}$$

As expected, the adiabatic enzyme profile $e^{(0)}$ agrees well with the enzyme profile obtained in a simulation with an enzyme-loaded static vesicle, i.e., a vesicle that cannot move and has a fixed position in space (Fig. S4a). For moving vesicles, $e^{(1)}$ captures the enzyme profile observed in the simulation (Fig. S4b).

##### B. Force and Velocity

The pressure created by a given enzyme profile in the vesicle can be computed via the ideal gas law (equivalent to the dilute limit of van't Hoff's law),

$$\Pi(r,t) = k_\mathrm{B}T\,e(r,t)\,, \tag{S66}$$

where $\Pi$ is the pressure and $e$ the enzyme concentration (number of enzymes per volume unit). The force exerted by the enzymes can be obtained by integrating the pressure over the whole membrane surface $S$,

$$\mathbf{F} = \oint_S \Pi(x)\mathbf{n}\,\mathrm{d}S = \int_V \nabla\Pi\,\mathrm{d}V = \mathbf{e_x}\;k_\mathrm{B}T\int_{-R}^{R}\pi(R^2-x^2)\partial_x e(x)\mathrm{d}x = \mathbf{e_x}2\pi kT\int_{-R}^{R}xe(x)\,\mathrm{d}x\,. \tag{S67}$$

In the second-to-last step, we again assumed that the enzyme profile is isotropic in the directions perpendicular to the direction of the gradient, i.e., isotropic in the $y$- and $z$-direction. Equivalently, we can express the force in terms of the dimensionless representation of the enzyme-profile,

$$\mathbf{F} = \mathbf{e_x}\,2\pi k_\mathrm{B}TR^2e_\mathrm{T}\int_{-1}^{1}\mathrm{d}\tilde{x}\;\tilde{x}\tilde{e}(\tilde{x}). \tag{S68}$$

Provided that the vesicle motion is overdamped and that the vesicle's friction is given by Stokes friction, velocity in $x$ direction resulting from the force is given by

$$v = \frac{F}{6\pi\eta R} = \frac{k_\mathrm{B}Te_\mathrm{T}R}{3\eta}\int_{-1}^{1}\mathrm{d}\tilde{x}\;\tilde{x}\tilde{e}(\tilde{x}). \tag{S69}$$

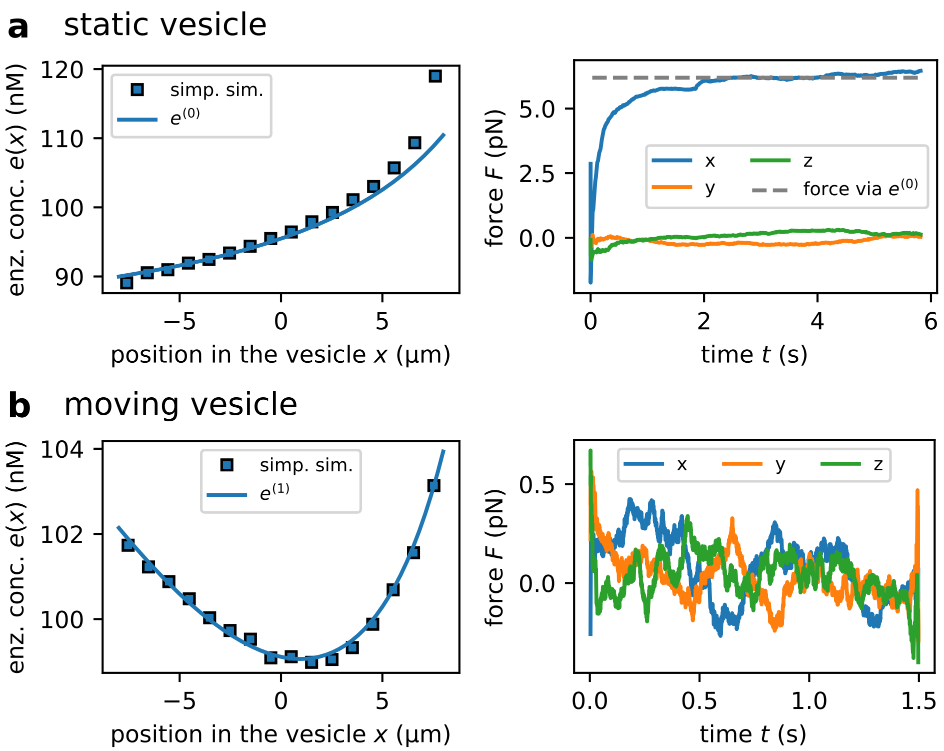

Figure S4. **Enzymes profile and forces in vesicles** Enzyme profile and forces observed in static (a) and freely moving vesicles (b). The data points are obtained using the simplified simulation, while the continuous curves show the analytically predicted enzyme profiles. For static vesicles, the enzyme profile observed in the simulation agrees well with the adiabatic enzyme profile $e^{(0)}$ (a); for moving vesicles, the simulated enzyme profile agrees with the analytical enzyme profile obtained to linear order in Péclet number (b). The simulation parameters are summarized in Table III.

For the adiabatic enzyme profile, $\tilde{e}^{(0)}(\tilde{x})$, we obtain the adiabatic velocity,

$$v^{(0)} = \frac{k_{\mathrm{B}} T e_{\mathrm{T}} R}{3\eta} \int_{-1}^{1} \mathrm{d}\tilde{x}\; \tilde{x}\tilde{e}^{(0)}(\tilde{x}) \tag{S70}$$

(the integral is evaluated explicitely in Sec. IV D and IV E). For the enzyme profile including contributions to linear order in Péclet number, we find the velocity,

$$\begin{aligned}
v^{(1)} &= \frac{k_{\mathrm{B}} T e_{\mathrm{T}} R}{3\eta} \int_{-1}^{1} \mathrm{d}\tilde{x}\; \tilde{x}\tilde{e}^{(1)}(\tilde{x}) && \text{(S71)}\\
&= \underbrace{\frac{kTe_{\mathrm{T}}R}{3\eta} \int_{-1}^{1} \mathrm{d}\tilde{x}\; \tilde{x}\tilde{e}^{(0)}(\tilde{x})}_{=v^{(0)}} + \mathrm{Pe}\,\mathcal{W}^{a} \underbrace{\frac{kTe_{\mathrm{T}}R}{3\eta} \int_{-1}^{1} \mathrm{d}\tilde{x}\; \tilde{x}\tilde{e}^{(0)}(\tilde{x})}_{=v^{(0)}} && \text{(S72)}\\
&\quad - \mathrm{Pe}\,\frac{kTe_{\mathrm{T}}R}{3\eta} \int_{-1}^{1} \mathrm{d}\tilde{x}\; \tilde{x}\tilde{e}^{(0)}(\tilde{x}) \int_{-1}^{\tilde{x}} \mathrm{d}\tilde{x}'\, \frac{1}{\tilde{D}_e(\tilde{x}')} && \text{(S73)}\\
&= v^{(0)} + \mathrm{Pe}\,\mathcal{W}^{a} v^{(0)} - \mathrm{Pe}\,\frac{kTe_{\mathrm{T}}R}{3\eta}\,\frac{4}{3}\,\frac{\int_{-1}^{1} \mathrm{d}\tilde{x}\; \tilde{x}\frac{1}{\tilde{D}_e(\tilde{x})} \int_{-1}^{\tilde{x}} \mathrm{d}\tilde{x}'\,\frac{1}{\tilde{D}_e(\tilde{x}')}}{\int_{-1}^{1} \mathrm{d}\tilde{x}\,\frac{1-\tilde{x}^2}{\tilde{D}_e(\tilde{x})}} && \text{(S74)}\\
&= v^{(0)} + \mathrm{Pe}\,\mathcal{W}^{a} v^{(0)} - \mathrm{Pe}\,\mathcal{W}^{b} v^{(0)}, && \text{(S75)}
\end{aligned}$$

where we defined a new Péclet number independent weight,

$$\mathcal{W}^b = \frac{1}{v^{(0)}}\frac{4}{3}\frac{kTe_{\rm tot}R}{3\eta}\frac{\int_{-1}^{1}\mathrm{d}\tilde{x}\,\frac{\tilde{x}}{\tilde{D}_e(\tilde{x})}\int_{-1}^{\tilde{x}}\mathrm{d}\tilde{x}'\frac{1}{\tilde{D}_e(\tilde{x}')}}{\int_{-1}^{1}\mathrm{d}\tilde{x}\frac{1-\tilde{x}^2}{\tilde{D}_e(\tilde{x})}} \tag{S76}$$

$$= \frac{3\eta}{kTe_{\rm tot}R}\frac{\int_{-1}^{1}\mathrm{d}\tilde{x}\,\frac{1-\tilde{x}^2}{\tilde{D}_e(\tilde{x})}}{\int_{-1}^{1}\mathrm{d}\tilde{x}\frac{\tilde{x}}{\tilde{D}_e(\tilde{x})}}\frac{kTe_{\rm tot}R}{3\eta}\frac{\int_{-1}^{1}\mathrm{d}\tilde{x}\,\frac{\tilde{x}}{\tilde{D}_e(\tilde{x})}\int_{-1}^{\tilde{x}}\mathrm{d}\tilde{x}'\frac{1}{\tilde{D}_e(\tilde{x}')}}{\int_{-1}^{1}\mathrm{d}\tilde{x}\frac{1-\tilde{x}^2}{\tilde{D}_e(\tilde{x})}} \tag{S77}$$

$$= \frac{\int_{-1}^{1}\mathrm{d}\tilde{x}\,\frac{\tilde{x}}{\tilde{D}_e(\tilde{x})}\int_{-1}^{\tilde{x}}\mathrm{d}\tilde{x}'\frac{1}{\tilde{D}_e(\tilde{x}')}}{\int_{-1}^{1}\mathrm{d}\tilde{x}\frac{\tilde{x}}{\tilde{D}_e(\tilde{x})}}\,. \tag{S78}$$

Summarizing the weight functions $\mathcal{W}^a$ and $\mathcal{W}^b$ into a single weight, $\mathcal{W} = \mathcal{W}^b - \mathcal{W}^a$, allows us to write the velocity as

$$v^{(1)} = v^{(0)} - \mathrm{Pe}\,\mathcal{W}v^{(0)} = v^{(0)} - \frac{vR}{D_{\rm e}^0}\mathcal{W}v^{(0)}, \tag{S79}$$

where we used the definition of Pe in the last step. Imposing a self-consistency constraint, i.e., assuming that the translation velocity, $v$, is well-approximated by the velocity obtained from the expansion to linear order in Péclet number, $v^{(1)}$, allows us to solve for $v = v^{(1)}$,

$$v^{(1)} = \frac{v^{(0)}}{1+\mathcal{W}\frac{v^{(0)}R}{D_{\rm e}^0}}. \tag{S80}$$

In the limit of small adiabatic Péclet number, $\mathrm{Pe}^{(0)} = v^{(0)}R/D_{\rm e}^0$, the translation velocity to linear order in Péclet number is well-approximated by the adiabatic velocity,

$$v^{(1)} \approx v^{(0)} \quad \text{if} \quad \mathcal{W}\mathrm{Pe}^{(0)} \ll 1. \tag{S81}$$

For large adiabatic velocity (large adiabatic Péclet number), the $v^{(1)}$ is independent of the adiabatic velocity, and only depends on $\mathcal{W}$,

$$v^{(1)} \approx \frac{D_{\rm e}^0}{R\mathcal{W}} \quad \text{if} \quad \mathcal{W}\mathrm{Pe}^{(0)} \gg 1. \tag{S82}$$

##### C. Effective Trajectory-Averaged Velocity

The translation velocity $v$ depends on the position of the vesicle in the system: As the vesicle moves, the substrate concentration within the vesicle changes, which affects the effective diffusion coefficient of the enzymes, and, consequently, the pressure and the ensuing translation velocity. In the derivation so far, we assumed that the center of the vesicle is placed in the middle of the system, $x = 0$, such that the vesicle "sees" the part of the substrate gradient between $x = -R$ and $x = R$ (the whole system extends from $x = -L/2$ to $x = L/2$). As the vesicle moves, the position of the center of the vesicle changes, $x = \xi$, and the relevant part of the substrate gradient is between $= \xi - R$ and $x = \xi + R$. We need to account for this by shifting the position at which the substrate gradient (and thus the effective diffusion coefficient) is evaluated, $x \to x + \xi$ (i.e., $\tilde{x} \to \tilde{x} + \xi/R$). With this shift, the full steady-state enzyme profile reads (analogous to Eq. (S57))

$$\tilde{e}(\tilde{x},\xi) = \frac{4}{3}\frac{\frac{1}{\tilde{D}_e(\tilde{x}+\xi/R)}\exp\left[-\frac{vR}{D_{\rm e}^0}\int_{-1}^{\tilde{x}}\mathrm{d}\tilde{x}'\frac{1}{\tilde{D}_e(\tilde{x}'+\xi/R)}\right]}{\int_{-1}^{1}\mathrm{d}\tilde{x}\,\frac{1-\tilde{x}^2}{\tilde{D}_e(\tilde{x}+\xi/R)}\exp\left[-\frac{vR}{D_{\rm e}^0}\int_{-1}^{\tilde{x}}\mathrm{d}\tilde{x}'\frac{1}{\tilde{D}_e(\tilde{x}'+\xi/R)}\right]}\,. \tag{S83}$$

Consequently, the velocity of the vesicle,

$$v(\xi) = \frac{k_{\rm B}Te_{\rm T}R}{3\eta}\int_{-1}^{1}\mathrm{d}\tilde{x}\,\tilde{x}\,\tilde{e}(\tilde{x},\xi), \tag{S84}$$

depends on the position of the vesicle and varies along its trajectory. The velocity observed in the simulation is the effective vesicle velocity $v^{\rm eff}$ averaged over the positions of the vesicle along its trajectory. To compute this effective velocity, we need to determine the distance by which the vesicle moves, $\Delta x(t) = x(t) - x(0)$, by integrating the equation of motion,

$$\frac{\mathrm{d}x(t)}{\mathrm{d}t} = v^{(1)}(x). \tag{S85}$$

Provided the change in the position of the vesicle is small enough, we can expand the velocity to linear order in $x$,

$$\frac{\mathrm{d}x(t)}{\mathrm{d}t} \approx v^{(1)}(x=0) + \left.\frac{\partial v^{(1)}(x)}{\partial x}\right|_{x=0} x. \tag{S86}$$

This differential equation can be solved analytically,

$$x(t) - x(0) = \frac{1}{t}\frac{v^{(1)}(x=0)}{\left.\frac{\partial v^{(1)}}{\partial x}\right|_{x=0}}\left[\exp\left(\left.\frac{\partial v}{\partial x}\right|_{x=0} t\right) - 1\right], \tag{S87}$$

leading to the following expression for the effective (trajectory-averaged) velocity,

$$v^{\mathrm{eff}}(t) = \frac{x(t) - x(0)}{t} = \frac{1}{t}\frac{v^{(1)}(x=0)}{\left.\frac{\partial v^{(1)}}{\partial x}\right|_{x=0}}\left[\exp\left(\left.\frac{\partial v}{\partial x}\right|_{x=0} t\right) - 1\right]. \tag{S88}$$

The velocity at position $x = 0$ to linear order in Péclet number, $v^{(1)}(x = 0)$, is computed via Eq. (S80). To evaluate $\partial_x v^{(1)}(x)|_{x=0}$, we make use of autodifferentiation. The theoretically predicted effective velocities are plotted in Fig. 5 as continuous curves, and agree well with the translation velocities observed in the simulation.

##### D. Substrate-Dependence of the Velocity

In this section we consider the substrate-dependence of the velocity. We assume that the substrate concentration equals zero on the right end of the system, while it attains a variable non-zero concentration $s_L$ on the left end of the system. As discussed before (see Section IV B), $v^{(1)}$ is well-approximated by the adiabatic velocity $v^{(0)}$ in the limit of small adiabatic Péclet number (i.e., $\mathcal{W}\,\mathrm{Pe}^{(0)} \ll 1$). Therefore, we need to determine the substrate-dependence of $v^{(0)}$ in order to understand the scaling of $v^{(1)}$ with the substrate in this limit.
Combining the definition of the adiabatic velocity (Eq. (S70)) with the adiabatic enzyme profile (Eq. (S58)), we find the following expression for the adiabatic velocity,

$$v^{(0)} = \frac{4k_{\mathrm{B}}Te_{\mathrm{T}}R}{9\eta}\frac{\int_{-1}^{1}\mathrm{d}\tilde{x}\,\frac{\tilde{x}}{\tilde{D}_e(\tilde{x})}}{\int_{-1}^{1}\mathrm{d}\tilde{x}\,\frac{1-\tilde{x}^2}{\tilde{D}_e(\tilde{x})}} = \frac{4k_{\mathrm{B}}Te_{\mathrm{T}}R}{9\eta}\frac{\mathcal{I}_1}{\mathcal{I}_2}, \tag{S89}$$

where we introduced the integrals,

$$\mathcal{I}_1 = \int_{-1}^{1}\mathrm{d}\tilde{x}\,\frac{\tilde{x}}{\tilde{D}_e(\tilde{x})}, \quad \text{and} \quad \mathcal{I}_2 = \int_{-1}^{1}\mathrm{d}\tilde{x}\,\frac{1-\tilde{x}^2}{\tilde{D}_e(\tilde{x})}. \tag{S90}$$

We recall that the dimensionless diffusion coefficient is a function of the position-dependent substrate concentration (Eq. (2)),

$$\tilde{D}_e(\tilde{x}) = 1 + \alpha\frac{s(\tilde{x})}{K_{\mathrm{M}} + s(\tilde{x})}, \tag{S91}$$

with the substrate concentration,

$$s(\tilde{x}) = \frac{s_L}{2} - \frac{s_L}{L}R\tilde{x}. \tag{S92}$$

As mentioned before, $s_L$ denotes the concentration on the left end of the system, and the concentration on the right end of the system is assumed to be zero. The linear position-dependence of the substrate allows us express the position $\tilde{x}$ as function of the concentration,

$$\tilde{x} = \frac{L}{2R} - \frac{L}{Rs_L}s \tag{S93}$$

such that we can re-parametrize the integrals with respect to $\tilde{x}$ (Eq. (S90)) as integrals with respect to $s$,

$$\mathcal{I}_1 = \int_{-1}^{1} \mathrm{d}\tilde{x}\ \frac{\tilde{x}}{\tilde{D}_e(\tilde{x})} = -\frac{L}{s_L R}\int_{\frac{s_L}{2}-\frac{s_L R}{L}}^{\frac{s_L}{2}+\frac{s_L R}{L}} \mathrm{d}s\ \frac{\frac{L}{2R}-\frac{L}{Rs_L}s}{\tilde{D}_e(s)}, \tag{S94}$$

$$\mathcal{I}_2 = \int_{-1}^{1} \mathrm{d}\tilde{x}\ \frac{1-\tilde{x}^2}{\tilde{D}_e(\tilde{x})} = -\frac{L}{s_L R}\int_{\frac{s_L}{2}-\frac{s_L R}{L}}^{\frac{s_L}{2}+\frac{s_L R}{L}} \mathrm{d}s\ \frac{1-\left(\frac{L}{2R}-\frac{L}{Rs_L}s\right)^2}{\tilde{D}_e(s)}. \tag{S95}$$

Unlike the integrals with respect to $\tilde{x}$, these integrals with respect to $s$ can be evaluated using a computer algebra system. To rationalize the scaling of $v^{(0)}$ with the substrate concentration at the left-end of the system, $s_L$, we use the expressions for the integrals $\mathcal{I}_1$ and $\mathcal{I}_2$ obtained via the computer algebra system to compute $v^{(0)}$, and expand $1/v^{(0)}$ in small concentrations,

$$\frac{1}{v^{(0)}} = \frac{9\eta}{4k_{\mathrm{B}}Te_{\mathrm{T}}R}\left(\frac{2K_{\mathrm{M}}L}{\alpha R}\frac{1}{s_L} + \frac{(2+\alpha)L}{\alpha R} + \frac{(1+\alpha)(5L^2-4(3+2\alpha)R^2)}{10(\alpha K_{\mathrm{M}}LR)}s_L\right) + \mathcal{O}(s_L^2). \tag{S96}$$

This result allows us to identify two regimes: (i) For small substrate concentrations, $1/v^{(0)}$ is proportional to $1/s_L$, implying that the velocity scales linearly with $v^{(0)} \sim s_L$, and (ii) for large substrate concentration, $1/v^{(0)}$ is proportional to $s_L$, implying that $v^{(0)} \sim 1/s_L$. The characteristic substrate concentration ${s_L^*}^{(0)}$ that separates the two regimes from each other is set by the condition that both contributions ($\sim s_L^{-1}$ and $\sim s_L$) need to contribute equally to the velocity,

$$\frac{2K_M L}{\alpha R {s_L^*}^{(0)}} = \frac{(1+\alpha)(5L^2-4(3+2\alpha)R^2){s_L^*}^{(0)}}{10(\alpha K_{\mathrm{M}}LR)}, \tag{S97}$$

which leads to the following expression for the characteristic substrate concentration,

$${s_L^*}^{a} = \frac{\sqrt{20}K_M L}{\sqrt{(1+\alpha)(5L^2-4(3+2\alpha)R^2)}}. \tag{S98}$$

Note that this concentration coincides with the position of the maximum, as can be verified by finding the concentration that solves $\partial_{s_L}(v^{(0)})^{-1} = 0$.

In the limit of large adiabatic Péclet number, $\mathcal{W}\,\mathrm{Pe}^{(0)} \gg 1$, $v^{(0)}$ does not approximate $v^{(1)}$ anymore. Instead, the velocity is given by

$$v^{(1)} \approx \frac{D_{\mathrm{e}}^0}{R\mathcal{W}}. \tag{S99}$$

To understand the substrate-dependence of the translation velocity in this limit, we need to evaluate $\mathcal{W}$ (Eq. (S64) and (S78)),

$$\mathcal{W} = \frac{\int_{-1}^{1}\mathrm{d}\tilde{x}\ \frac{\tilde{x}}{\tilde{D}_e(\tilde{x})}\int_{-1}^{\tilde{x}}\mathrm{d}\tilde{x}'\frac{1}{\tilde{D}_e(\tilde{x}')}}{\int_{-1}^{1}\mathrm{d}\tilde{x}\frac{\tilde{x}}{\tilde{D}_e(\tilde{x})}} - \frac{\int_{-1}^{1}\mathrm{d}\tilde{x}\ \frac{1-\tilde{x}^2}{\tilde{D}_e(\tilde{x})}\int_{-1}^{\tilde{x}}\mathrm{d}\tilde{x}'\frac{1}{\tilde{D}_e(\tilde{x}')}}{\int_{-1}^{1}\mathrm{d}\tilde{x}\frac{1-\tilde{x}^2}{\tilde{D}_e(\tilde{x})}} = \frac{\mathcal{I}_3}{\mathcal{I}_1} - \frac{\mathcal{I}_4}{\mathcal{I}_2}, \tag{S100}$$

where we re-used the integrals $\mathcal{I}_1$ and $\mathcal{I}_2$ introduced previously, and defined,

$$\mathcal{I}_0(\tilde{x}) = \int_{-1}^{\tilde{x}}\mathrm{d}\tilde{x}'\frac{1}{\tilde{D}_e(\tilde{x}')},\quad \mathcal{I}_3 = \int_{-1}^{1}\mathrm{d}\tilde{x}\ \frac{\tilde{x}}{\tilde{D}_e(\tilde{x})}\mathcal{I}_0(\tilde{x}),\quad \mathcal{I}_4 = \int_{-1}^{1}\mathrm{d}\tilde{x}\ \frac{1-\tilde{x}^2}{\tilde{D}_e(\tilde{x})}\mathcal{I}_0(\tilde{x}) \tag{S101}$$

Expressing the dimensionless position $\tilde{x}$ in terms of the substrate concentration $s$ (Eq. (S93)), allows us to identify the following representation of the integrals,

$$\mathcal{I}_0(s) = -\frac{L}{s_L R}\int_{\frac{s_L}{2}-\frac{s_L R}{L}}^{s}\mathrm{d}s'\frac{1}{\tilde{D}_e(s)}, \tag{S102}$$

$$\mathcal{I}_3 = -\frac{L}{s_L R}\int_{\frac{s_L}{2}-\frac{s_L R}{L}}^{\frac{s_L}{2}+\frac{s_L R}{L}}\mathrm{d}s\ \frac{\mathcal{I}_0(s)\left(\frac{L}{2R}-\frac{L}{Rs_L}s\right)}{\tilde{D}_e(s)}, \tag{S103}$$

$$\mathcal{I}_4 = -\frac{L}{s_L R}\int_{\frac{s_L}{2}-\frac{s_L R}{L}}^{\frac{s_L}{2}+\frac{s_L R}{L}}\mathrm{d}s\ \frac{\mathcal{I}_0(s)\left[1-\left(\frac{L}{2R}-\frac{L}{Rs_L}s\right)^2\right]}{\tilde{D}_e(s)}. \tag{S104}$$

Using a computer algebra system to evaluate these integrals allows us to find an analytical expression for $\mathcal{W}$. Expanding in small $s_L$, we obtain the following expression for $\mathcal{W}$,

$$\mathcal{W} = \frac{K_\mathrm{M} L}{\alpha R}\frac{1}{s_L} + \frac{L}{\alpha R} + \frac{5L^2 - 4(3+2\alpha-\alpha^2)R^2}{20\alpha K_\mathrm{M} L R} s_L + \mathcal{O}(s_L^2). \tag{S105}$$

Just as previously, this implies that there are two regimes: (i) For small $s_L$, the first term in Eq. (S105) dominates, such that $\mathcal{W} \sim s_L^{-1}$. As the velocity is inversely proportional to $\mathcal{W}$, the velocity scales linearly in substrate concentration, $v^{(1)} \sim s_L$ for small substrate concentrations. (ii) For large $s_L$, the third term in Eq. (S105) is contributes most significantly, such that the velocity decays as $v^{(1)} \sim s_L^{-1}$. We can again identify the characteristic substrate concentration for the transition between the regimes (i.e., the concentration at which both terms contribute equally to the velocity),

$$s_L^{*\,b} = \frac{\sqrt{20} K_M L}{\sqrt{5L^2 - 4(3+2\alpha-\alpha^2)R^2}} \tag{S106}$$

Note that this characteristic substrate concentration $s_L^{*\,b}$ can differ from the characteristic concentration obtained for the adiabatic velocity, $s_L^{*\,a}$. In summary, we find that

$$v^{(1)} \sim \begin{cases} s_L^{-1} & \text{if} \quad \mathcal{W}\mathrm{Pe}^{(0)} \ll 1 \quad \text{and} \quad s_L \ll s_L^{*\,a}, \\ s_L & \text{if} \quad \mathcal{W}\mathrm{Pe}^{(0)} \ll 1 \quad \text{and} \quad s_L \gg s_L^{*\,a}, \\ s_L^{-1} & \text{if} \quad \mathcal{W}\mathrm{Pe}^{(0)} \gg 1 \quad \text{and} \quad s_L \ll s_L^{*\,b}, \\ s_L & \text{if} \quad \mathcal{W}\mathrm{Pe}^{(0)} \gg 1 \quad \text{and} \quad s_L \gg s_L^{*\,b}. \end{cases} \tag{S107}$$

For a vesicle moving in a medium of high viscosity (i.e., $\eta = 1\,\mathrm{Pa\,s}$), the velocity of translation is well-approximated by the adiabatic velocity $v^{(0)}$ over the entire range of studied substrate concentrations (Fig. S5A-B). Thus, the first two cases in Eq. (S107) apply: The velocity reaches a maximum at $s_L^{*\,a}$. For concentrations smaller than $s_L^{*\,a}$, the velocity increases linearly with $s_L$, while it decreases linearly with $s_L^{-1}$ for concentrations larger than $s_L^{*\,a}$ (Fig. S5C). Fig. S5 shows the velocity of a vesicle placed at $x = 0$ (no trajectory averaging!), the trajectory-averaged effective velocity is plotted in Fig. S11A. Given the small velocities (and consequently small displacement), the effecitve velocity and the velocity at $x = 0$ are almost identical.
If the vesicle moves in a medium of water-like viscosity (i.e., $\eta = 1\,\mathrm{mPa\,s}$), the velocity of translation is well-approximated by the $D_\mathrm{e}^0 / (\mathcal{W}R)$ over the entire range of studied substrate concentrations (Fig. S6A-B). Thus, the last two cases in Eq. (S107) apply: The velocity reaches a maximum at $s_L^{*\,b}$. For concentrations smaller than $s_L^{*\,b}$, the velocity increases linearly with $s_L$, while it decreases linearly with $s_L^{-1}$ for concentrations larger than $s_L^{*\,b}$ (Fig. S6C). Again, Fig. S6 depicts the translation velocity for a vesicle at $x = 0$, the respective effective velocity is plotted in Fig. 5A.

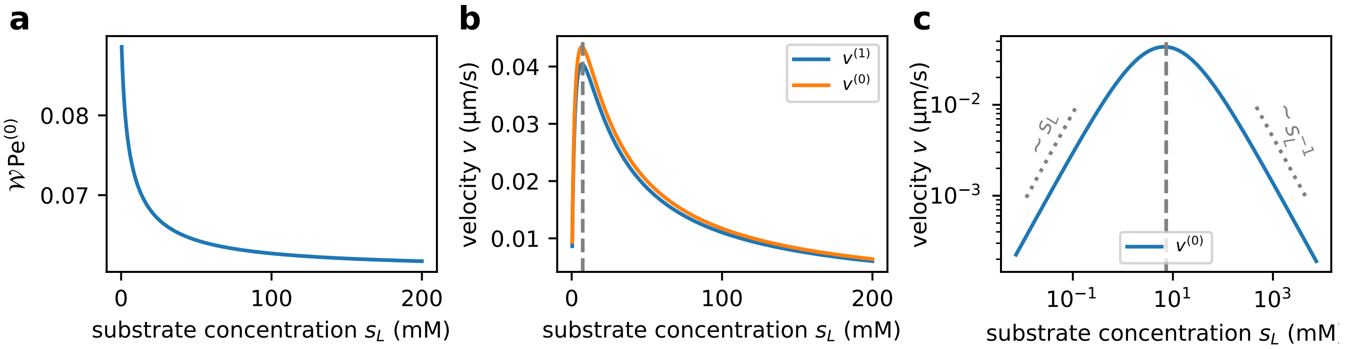

Figure S5. **Substrate-dependence of the translation velocity for high viscosity, $\eta = 1\,\mathrm{Pa\,s}$ (without trajectory averaging of the velocity)**. a) The order parameter $\mathcal{W}\mathrm{Pe}^{(0)}$ is small over the entire studied range of parameters, implying that the translation velocity $v^{(1)}$ is well-approximated by the adiabatic velocity $v^{(0)}$ (see panel B). b) The translation velocity exhibits a non-monotoneous dependence on the substrate concentration, reaching a maximum at intermediary substrate concentration, $s_L = s_L^{*\,a}$ (see grey vertical dashed line). c) For small substrate concentration, $v^{(0)}$ is proportional to $s_L$, while for large substrate concentration $v^{(0)} \sim s_l^{-1}$ (see power laws plotted as dashed lines). In all panels, the velocities are computed using the system parameters summarized in Table III. The viscosity of the medium is similar to that of water, $\eta = 1\,\mathrm{mPa\,s}$.

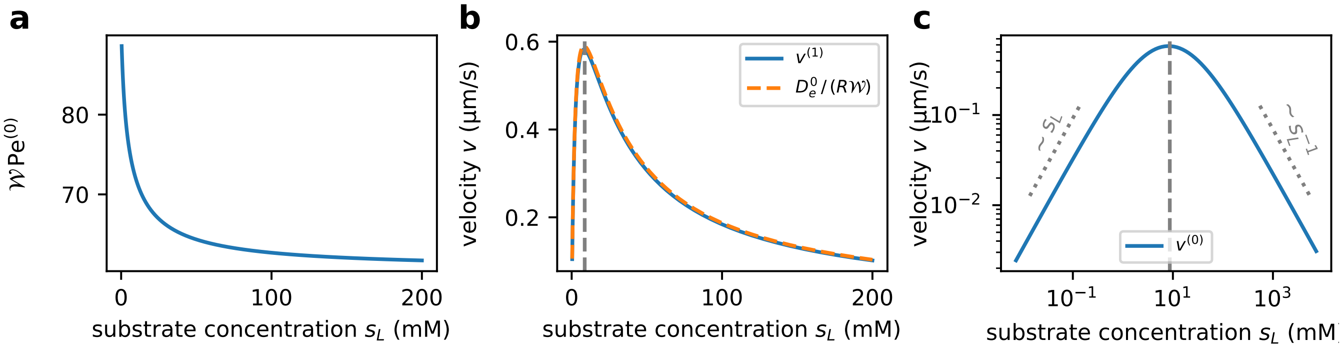

Figure S6. **Substrate-dependence of the translation velocity for small viscosity,** $\eta = 1\,\mathrm{mPa\,s}$ **(without trajectory averaging of the velocity).** a) The order parameter $\mathcal{W}\mathrm{Pe}^{(0)}$ is much larger than 1 over the entire studied range of parameters, implying that the translation velocity $v^{(1)}$ is well-approximated by the $D_\mathrm{e}^0\,/\,(R\mathcal{W})$ (see panel B). b) The translation velocity exhibits a non-monotoneous dependence on the substrate concentration, reaching a maximum at intermediary substrate concentration, $s_L = s_L^{*\,b}$ (see grey vertical dashed line). c) For small substrate concentration, $v^{(0)}$ is proportional to $s_L$, while for large substrate concentration $v^{(0)} \sim s_l^{-1}$ (see power laws plotted as dashed lines). All parameters (except for the viscosity) are identical to the parameters summarized in Table III.

##### E. Radius-Dependence of the Velocity

Similarly as in the last section, we analyze the radius-dependence of $v^{(1)}$ in the limit of small and high adiabatic Péclet number separately. In the limit of small adiabatic Péclet number, $v^{(1)}$ is well-approximated by $v^{(0)}$. We start by analyzing the radius-dependence of $v^{(0)}$: We use the representation of the adiabatic velocity in terms of the integrals $\mathcal{I}_1$ and $\mathcal{I}_2$ introduced in the last section (Eqs. (S89) and (S90)), and expand the expression for $v^{(0)}$ (obtained via a computer algebra system) in small vesicle radius $R$,

$$v^{(0)} = \frac{4k_\mathrm{B}Te_\mathrm{T}}{9\eta}\frac{2\alpha K_\mathrm{M}s_LR^2}{L(2K_\mathrm{M}+s_L)(2K_\mathrm{M}+(1+\alpha)s_L)} + \mathcal{O}(R^3). \tag{S108}$$

Hence, in the limit $\mathcal{W}\mathrm{Pe}^{(0)} \ll 1$, $v^{(1)}$ depends quadratically on the vesicle radius, in the same way as $v^{(0)}$. However, in the limit $\mathcal{W}\,\mathrm{Pe}^{(0)} \gg 1$, the velocity is given by $v^{(1)} \approx D_\mathrm{e}^0\,/\,(R\mathcal{W})$. To understand the radius dependence of the translation velocity in this limit, we express $\mathcal{W}$ in terms of the integrals $\mathcal{I}_0(s)$, $\mathcal{I}_3$ and $\mathcal{I}_4$ introduced previously (Eq. (S101)). Expanding $\mathcal{W}$ in small $R$, we obtain the following expression for $\mathcal{W}$,

$$\mathcal{W} \approx \mathcal{W}_{-1}R^{-1} + \mathcal{W}_1R + \mathcal{W}_3R^3, \tag{S109}$$

with radius-independent constants,

$$\mathcal{W}_{-1} = \frac{L(2K_M+s_L)^2}{4\alpha K_M s_L}, \tag{S110}$$

$$\mathcal{W}_1 = \frac{(1+\alpha)\;s_L\;(2K_M+s_L)\;[2(\alpha-3)K_M-3(1+\alpha)s_L]}{5\alpha k_M L(2K_M+(1+\alpha)s_L)^2}. \tag{S111}$$

This representation of $\mathcal{W}$ implies the following radius-dependence for the velocity,

$$v^{(1)} \approx \frac{D_\mathrm{e}^0}{\mathcal{W}_{-1}+\mathcal{W}_1R^2+\mathcal{W}_3R^4}. \tag{S112}$$

In addition to the scaling of $v^{(1)}$ for large radii, the expression for $\mathcal{W}$ (Eq. (S109)) also allows us to derive an analytical expression for the characteristic radius up to which the velocity $v^{(1)}$ is well-approximated by the adiabatic velocity. To this end, we use the criterion,

$$\mathcal{W}\,\mathrm{Pe}^{(0)} = \frac{\mathcal{W}(R^*)v^{(0)}(R^*)R^*}{D_\mathrm{e}^0} = 1, \tag{S113}$$

as well as the radius-dependent expression for $\mathcal{W} \approx \mathcal{W}_{-1}/R$ and $v^{(0)}$ (Eq. (S108)),

$$\frac{1}{D_\mathrm{e}^0}\frac{L(2K_\mathrm{M}+s_L)^2}{4\alpha K_\mathrm{M}s_L}\frac{4k_\mathrm{B}Te_\mathrm{T}}{9\eta}\frac{2\alpha K_\mathrm{M}s_L(R^*)^2}{L(2K_\mathrm{M}+s_L)(2K_\mathrm{M}+(1+\alpha)s_L)} = 1 \tag{S114}$$

$$\frac{2K_\mathrm{M}+s_L}{D_\mathrm{e}^0}\frac{2k_\mathrm{B}Te_\mathrm{T}}{9\eta}\frac{(R^*)^2}{(2K_\mathrm{M}+(1+\alpha)s_L)} = 1, \tag{S115}$$

which leads to the following expression for the characteristic radius,

$$R^* = \sqrt{\frac{9\eta D_\mathrm{e}^0\,(2K_\mathrm{M}+(1+\alpha)s_L)}{2k_\mathrm{B}Te_\mathrm{T}\,(2K_\mathrm{M}+s_L)}}. \tag{S116}$$

In summary, we find,

$$v^{(1)} \approx \begin{cases} v^{(0)} & \sim R^2 & \text{for} \quad R < R^*, \\ \dfrac{D_\mathrm{e}^0}{R\mathcal{W}} & \sim (\mathcal{W}_{-1}+\mathcal{W}_1R^2+\mathcal{W}_3R^4)^{-1} & \text{for} \quad R > R^*. \end{cases} \tag{S117}$$

For a vesicle moving in a medium of high viscosity (i.e., $\eta = 1\,\mathrm{Pa\,s}$), $R^* \approx 30\,\mathrm{\mu m}$ for the investigated parameters, implying that the velocity scales quadratically with the radius for all studied radii, as shown in Fig. S7 (adiabatic velocity without trajectory-averaging) and Fig. S11 (effective velocity with trajectory-averaging).
If the vesicle moves in a medium of water-like viscosity (i.e., $\eta = 1\,\mathrm{mPa\,s}$), the characteristic radius equals $R^* \approx 1\,\mathrm{\mu m}$. Consequently, we observe both regimes introduced in Eq. (S117), as shown in Fig. S8B or Fig. 5b (see orange curve for $v^{(0)}$ and green curve for $D_\mathrm{e}^0/(R\mathcal{W})$ as well as the vertical dashed line representing $R^*$). Note that the representation in Fig. 5 shows the trajectory-averaged velocity computed based on $v^{(1)}$, while Fig. S8B shows the bare velocity $v^{(1)}$ without trajectory averaging. Moreover, we find that Eq. (S112) is indeed a good approximation of $D_\mathrm{e}^0/(R\mathcal{W})$ (Fig. S8C).

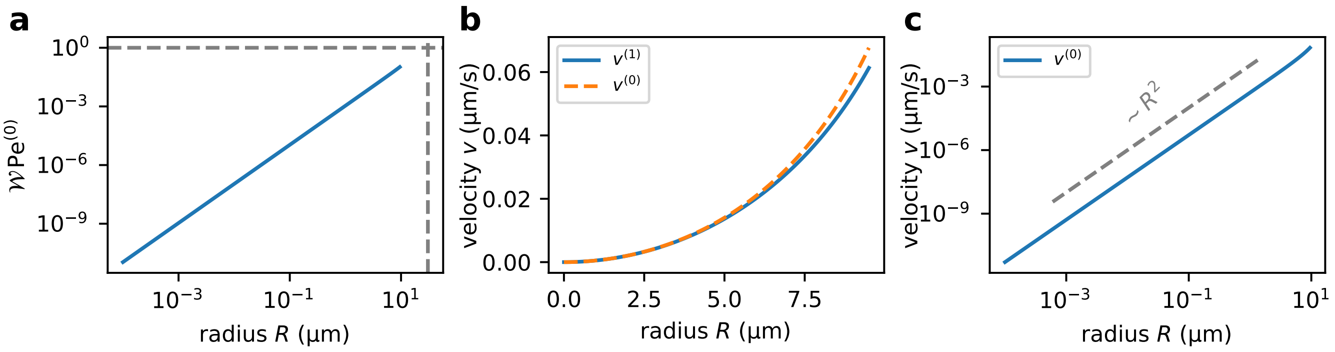

Figure S7. **Radius-dependence of the vesicle velocity for high viscosity, $\eta = 1\,\mathrm{Pa\,s}$ (without trajectory averaging).** a) The order parameter $\mathcal{W}\,\mathrm{Pe}^{(0)}$ is smaller than 1 over the entire range of studied radii. The characteristic radius $R^*$ (see vertical dashed line) is larger than the largest radius to be analyzed. b) Over the entire range of studied radii, $v^{(1)}$ is well-approximated by $v^{(0)}$. c) The adiabatic velocity $v^{(0)}$ (and thus $v^{(1)}$) scales quadratically with the vesicle radius $R$. All parameters are identical to the ones listed in Table III with a fixed substrate concentration of $s_L = 10\,\mathrm{mM}$.

##### F. Dependence of the Velocity on the Diffusion Coefficient

To understand the dependence of the translation velocity $v^{(1)}$ on the diffusion coefficient, we analyze how the adiabatic velocity $v^{(0)}$ and $D_\mathrm{e}^0\,/\,(R\mathcal{W})$ depend on $D_\mathrm{e}^0$:

- The adiabatic velocity $v^{(0)}$ is independent of $D_\mathrm{e}^0$, since the integrals $\mathcal{I}_1$ and $I_\in$ appearing in $v^{(0)}$ (Eq. (S89)) only depend on the *dimensionless* diffusion coeffcient $\tilde{D}_e(\tilde{x})$ (and therefore not on $D_\mathrm{e}^0$).
- $D_\mathrm{e}^0\,/\,(R\mathcal{W})$ depends linearly on $D_\mathrm{e}^0$, as the integrals appearing in $\mathcal{W}$ (Eq. (S100)) are independent of $D_\mathrm{e}^0$.

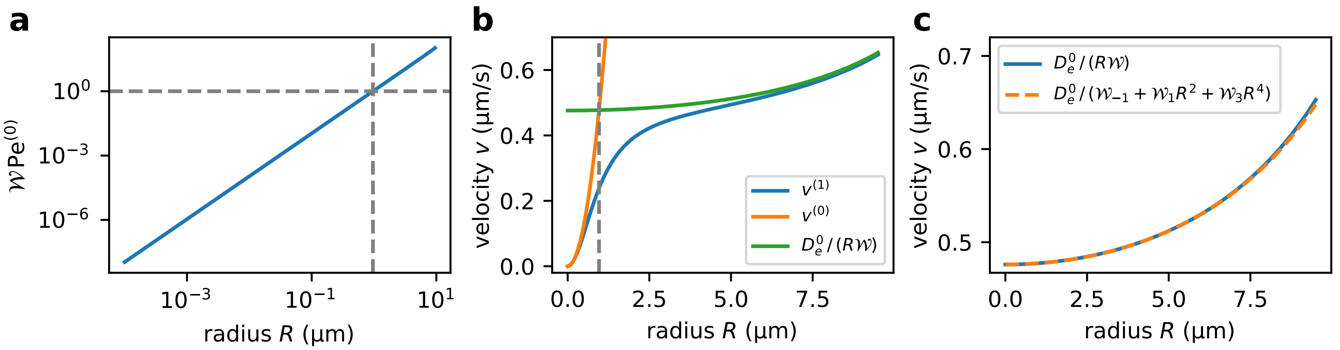

Figure S8. **Radius-dependence of the vesicle velocity for small viscosity, $\eta = 1\,\mathrm{mPa\,s}$ (without trajectory averaging).** a) The order parameter $\mathcal{W}\,\mathrm{Pe}^{(0)}$ is smaller than 1 for $R < R^*$, and larger than 1 for $R > R^*$. The characteristic radius $R^*$ (vertical dashed line) is set by the condition $\mathcal{W}\,\mathrm{Pe}^{(0)} = 1$ (horizontal dashed line). b) For small radii, $R < R^*$, $v^{(1)}$ is well-approximated by $v^{(0)}$, while $v^{(1)} \approx D_\mathrm{e}^0/(R\mathcal{W})$ for large radii, $R > R^*$. The characteristic radius $R^*$ is shown as vertical dashed line. c) $D_\mathrm{e}^0/(\mathcal{W}_{-1} + \mathcal{W}_1 R + \mathcal{W}_3 R^4)$ is a good approximation for $D_\mathrm{e}^0/(R\mathcal{W})$ over the entire studied range of radii. All parameters (except for the viscosity) are identical to the ones listed in Table III.

In summary, we find,

$$v^{(1)} \sim \begin{cases} 1 & \text{if} \quad \mathcal{W}\mathrm{Pe}^{(0)} \ll 1, \\ D_\mathrm{e}^0 & \text{if} \quad \mathcal{W}\mathrm{Pe}^{(0)} \gg 1. \end{cases} \tag{S118}$$

For a vesicle that moves in a medium of high viscosity (i.e., $\eta = 1\,\mathrm{Pa\,s}$), the translation velocity $v^{(1)}$ is set by $D_\mathrm{e}^0\,/\,(\mathcal{W}R)$ as long as the diffusion coefficient is small, such that the velocity increases linearly with $D_\mathrm{e}^0$ (see green dashed line in Fig. S9b). As the diffusion coefficient $D_\mathrm{e}^0$ increases, the dimensionless order parameter $\mathcal{W}\,\mathrm{Pe}^{(0)}$ decreases, implying that $v^{(1)}$ is well-approximated by $v^{(0)}$. Consequently, the velocity $v^{(1)}$ approaches the adiabatic velocity $v^{(0)}$ asymptotically, and $v^{(1)}$ is independent of $D_\mathrm{e}^0$ in the limit of high diffusion coefficients.

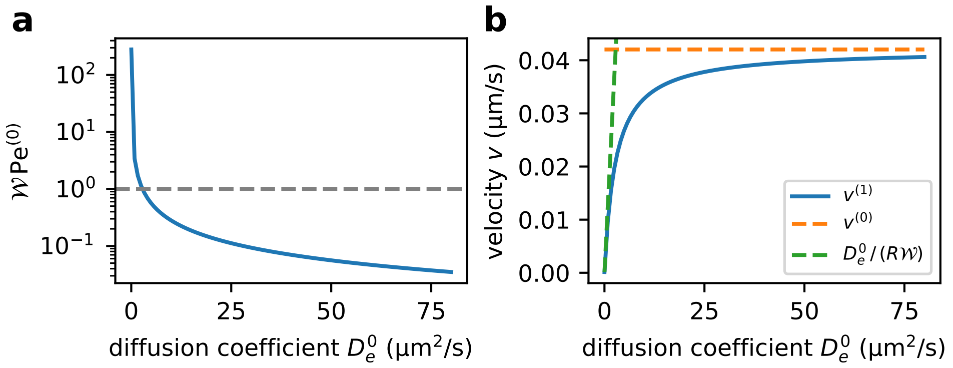

Figure S9. **Dependence of the vesicle velocity on the diffusion coefficient for high viscosity, $\eta = 1\,\mathrm{Pa\,s}$ (without trajectory averaging).** a) The order parameter $\mathcal{W}\,\mathrm{Pe}^{(0)}$ is smaller than 1 for small diffusion coefficient, but falls below 1 for larger $D_\mathrm{e}^0$. b) For small diffusion coefficient, $v^{(1)}$ is proportional to $D_\mathrm{e}^0$ as it is set by $D_\mathrm{e}^0/(R\,\mathcal{W})$ (green dashed line). For sufficiently large $D_\mathrm{e}^0$, $v^{(1)}$ approaches the $D_\mathrm{e}^0$-independent adiabatic velocity $v^{(0)}$ (orange dashed line). All parameters are identical to the ones listed in Table III with a fixed substrate concentration of $s_L = 10\,\mathrm{mM}$.

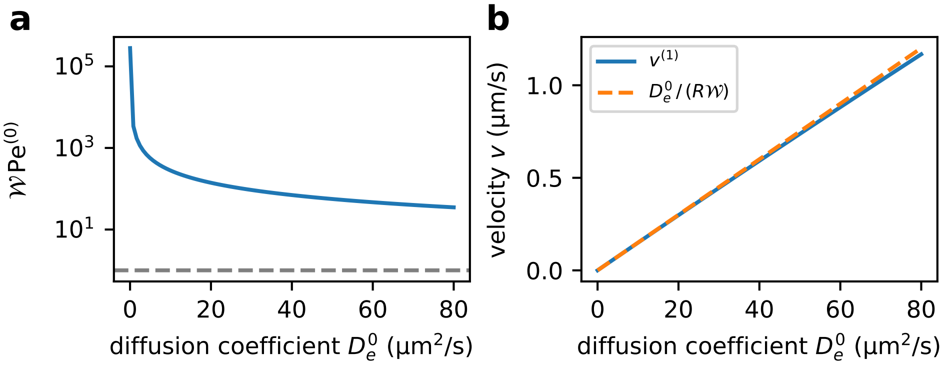

Figure S10. **Dependence of the vesicle velocity on the diffusion coefficient for small viscosity,** $\eta = 1\,\mathrm{mPa\,s}$ **(without trajectory averaging).** a) The order parameter $\mathcal{W}\,\mathrm{Pe}^{(0)}$ is larger than 1 over the entire range of diffusion coefficients $D_\mathrm{e}^0$. b) The velocity $v^{(1)}$ is proportional to $D_\mathrm{e}^0$ as it is set by $D_\mathrm{e}^0/(R\,\mathcal{W})$ (green dashed line). All parameters (except for the viscosity) are identical to the parameters listed in Table III.

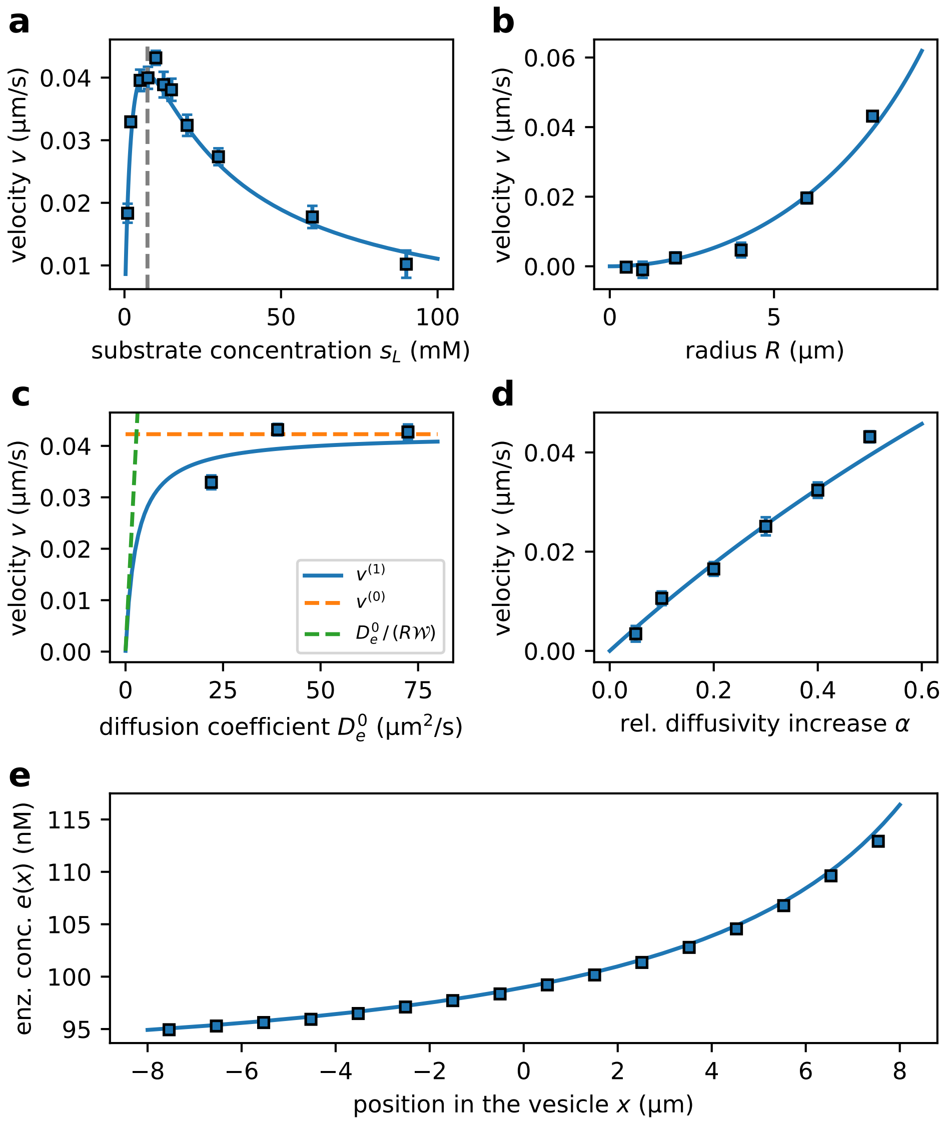

Figure S11. **Parameter-dependence of the vesicle for high viscosity** a) Vesicle velocity $v$ depends non-monotonously on the sustrate gradient, which is set by the substrate concentration on the left edge of the system, $s_L$. b) Vesicle velocity $v$ increases quadratically as a function of vesicle radius $R$. c) Vesicle velocity $v$ increases with the diffusion coefficient $D_\mathrm{e}^0$, and approaches the adiabatic velocity $v^{(0)}$ asymptotically in the limit of high $D_\mathrm{e}^0$. d) Vesicle velocity $v$ increases with the strength of enhanced diffusion, $\alpha$. e) Steady-state enzyme concentration profile. All panels show the translation velocity in a medium with viscosity that is a factor 1000 higher than that of water, $\eta = 1\,\mathrm{Pa\,s}$. All other parameters are summarized in Table III.

[1] R. R. Mayrand and D. G. Levitt, J. Gen. Physiol. **81**, 221 (1983).
[2] J. Winkelmann, "Diffusion coefficient of urea in water: Datasheet from physical chemistry · volume 15b2: "diffusion in gases, liquids and electrolytes" in springermaterials," (2018).
[3] B. Yang, "Transport characteristics of urea transporter-b," in *Urea Transporters* (Springer Netherlands, 2014) pp. 127–135.
[4] L. Van de Cauter, F. Fanalista, L. van Buren, N. De Franceschi, E. Godino, S. Bouw, C. Danelon, C. Dekker, G. H. Koenderink, and K. A. Ganzinger, ACS Synth. Biol. **10**, 1690 (2021).
[5] L. Song, M. R. Hobaugh, C. Shustak, S. Cheley, H. Bayley, and J. E. Gouaux, Science **274**, 1859 (1996).
[6] L. Y. Huang, W. A. Catterall, and G. Ehrenstein, J. Gen. Physiol. **71**, 397 (1978).
[7] H. R. Vutukuri, M. Hoore, C. Abaurrea-Velasco, L. van Buren, A. Dutto, T. Auth, D. A. Fedosov, G. Gompper, and J. Vermant, Nature **586**, 52 (2020).
[8] N. Kučerka, J. F. Nagle, J. N. Sachs, S. E. Feller, J. Pencer, A. Jackson, and J. Katsaras, Biophys. J. **95**, 2356 (2008).
[9] N. Kucerka, S. Tristram-Nagle, and J. Nagle, J. Membr. Biol. **208**, 193 (2006).
[10] N. Kucerka, S. Tristram-Nagle, and J. Nagle, Biophys. J. **90**, L83 (2006).
[11] P. Peterlin, G. Jaklič, and T. Pisanski, Meas. Sci. Technol. **20**, 055801 (2009).
[12] H. Saito and W. Shinoda, J. Phys. Chem. B **115**, 15241 (2011).
[13] W. G. Hill, R. L. Rivers, and M. L. Zeidel, J. Gen. Physiol. **114**, 405 (1999).
[14] M. Palaiokostas, W. Ding, G. Shahane, and M. Orsi, Soft Matter **14**, 8496 (2018).
[15] D. Cohen-Steiner and F. Da, Visual Comput. **20**, 4 (2004).
[16] T. K. F. Da and D. Cohen-Steiner, The Computational Geometry Algorithms Library (CGAL) User and Reference Manual (2023).
[17] K. Itô, Proceedings of the Imperial Academy of Japan **20**, 519 (1944).
[18] N. G. van Kampen, *Stochastic Processes in Physics in Chemistry* (Elsevier, 2007).
[19] J. Agudo-Canalejo, P. Illien, and R. Golestanian, Nano Lett. **18**, 2711 (2018).
[20] C. Riedel, R. Gabizon, M. C. A. Wilson, K. Hamadani, K. Tsekouras, S. Marqusee, S. Presse, and C. Bustamante, Nature **517**, 227 (2015).
[21] H. Noguchi and G. Gompper, Phys. Rev. E **72**, 011901 (2005).
[22] P. Iyer, G. Gompper, and D. A. Fedosov, Soft Matter **18**, 6868 (2022).
[23] A.-Y. Jee, T. Tlusty, and S. Granick, Proc. Natl. Acad. Sci. USA **117**, 29435 (2020).
